# The impact of common variants on gene expression in the human brain: from RNA to protein to schizophrenia risk

**DOI:** 10.1101/2023.06.04.543603

**Authors:** Qiuman Liang, Yi Jiang, Annie W. Shieh, Dan Zhou, Rui Chen, Feiran Wang, Meng Xu, Mingming Niu, Xusheng Wang, Dalila Pinto, Yue Wang, Lijun Cheng, Ramu Vadukapuram, Chunling Zhang, Kay Grennan, Gina Giase, The PsychENCODE Consortium, Kevin P White, Junmin Peng, Bingshan Li, Chunyu Liu, Chao Chen, Sidney H. Wang

## Abstract

**Background:** The impact of genetic variants on gene expression has been intensely studied at the transcription level, yielding invaluable insights into the association between genes and the risk of complex disorders, such as schizophrenia (SCZ). However, the downstream impact of these variants and the molecular mechanisms connecting transcription variation to disease risk are not well understood.

**Results:** We quantitated ribosome occupancy in prefrontal cortex samples of the BrainGVEX cohort. Together with transcriptomics and proteomics data from the same cohort, we performed cis-Quantitative Trait Locus (QTL) mapping and identified, at 10% false discovery rate, 3,253 expression QTLs (eQTLs), 1,344 ribosome occupancy QTLs (rQTLs), and 657 protein QTLs (pQTLs) out of 7,458 genes from 185 samples. Of the eQTLs identified, only 34% have their effects propagated to the protein level. Further analysis on the effect size of prefrontal cortex eQTLs identified from an independent dataset clearly replicated the post-transcriptional attenuation of eQTL effects. We identified omics-specific QTLs and investigated their potential in driving disease risks. Using both a variant-based approach and a gene-based approach, we identified genes containing expression-specific QTLs (esQTLs), ribosome-occupancy-specific QTLs (rsQTLs), and protein-specific QTLs (psQTLs). Among the variant-based omics-specific QTL, 38 showed strong colocalization with brain associated disorder GWAS signals, 29 of them are esQTLs. From the gene-based approach, we found 11 brain associated disorder risk genes that are driven predominantly by omics-specific QTL, all of them are driven by variants impacting transcriptional regulation. To take a complementary approach to further investigate the functional relevance of genes driven predominantly by attenuated eQTL signals, we identified SCZ risk genes using each omics independently and then investigated the omics-specificity of the driver regulatory process for each risk gene. Using S-PrediXcan we identified 74 SCZ risk genes across the three omics, 30% of which were novel, and 67% of these risk genes were confirmed to be causal in a MR-Egger test. Notably, 52 out of the 74 risk genes were identified using eQTL data and 68% of these SCZ-risk-gene-driving eQTLs show little to no evidence of driving corresponding variations at the protein level.

**Conclusion:** The effect of eQTLs on gene expression in the prefrontal cortex is commonly attenuated post-transcriptionally. Many of the attenuated eQTLs still correlate with GWAS signals of brain associated complex disorders, indicating the possibility that these eQTL variants drive disease risk through mechanisms other than regulating protein expression level. Further investigation is needed to elucidate the mechanistic link between attenuated eQTLs and brain associated complex disorders.

## Introduction

Complex diseases such as neuropsychiatric disorders are multi-factorial with genetic components (*1*, *2*). Large scale Genome-Wide Association Studies (GWAS) have uncovered thousands of disease associated loci, signaling a promising era ahead for causal variant identification (*3*). However, efforts in fine mapping these disease risk loci often narrowed down the underlying causal signals to non-coding regions of the genome (*4–6*). Regulatory variants in such non-coding regions are therefore the prime candidates for driving the genetic risk of disease etiology. Consequently, integrating gene expression information to pinpoint causal variants or to identify risk genes has become a staple of genetic studies of complex diseases (*7*), with multiple consortia efforts facilitating large scale gene expression profiling and regulatory element mapping (*8–10*). Many powerful methods, such as *coloc*, PrediXcan, SMR/HEIDI, to name a few, have also been developed to leverage gene expression information for fine mapping GWAS signals or for identifying underlying risk genes (*11–13*).

Brain associated complex disorders, such as neuropsychiatric disorders, are a group of complex diseases that can be highly heritable (*14*). For example, schizophrenia (SCZ), a psychiatric disorder affecting ∼1% of the world-wide population (*15*), has a heritability estimate ranged between 60% to 80%, indicating a strong genetic component (*16*, *17*). Accordingly, recent SCZ GWAS study reported by Trubetskoy et al. identified 287 significant risk loci and prioritized 120 risk genes using functional genomics (*18*). The use of RNA-Seq data from the brain was instrumental for risk gene prioritization by Trubetskoy et al., however, information from downstream gene regulation processes, such as translation rate and protein abundance, was either not utilized or unavailable.

Measuring transcriptional changes as a proxy for gene activity has a long history in molecular biology (*19*). In the context of human genetics, buffering of downstream effects of genetic variants impacting gene expression (i.e. an eQTL) has been shown to be prevalent (*20*). In addition, QTLs specific to protein level have also been reported (*20–22*). Together, these observations indicated the importance and potential benefits of including downstream omics types, such as proteomics data, as information sources for fine mapping disease regulatory variants. Indeed, recent studies using multi-omics approaches have demonstrated increased power for risk gene identification among other benefits (*23*, *24*). Of note, our recent work on genetic variants associated with protein expression level in prefrontal cortex of the human brain indicated the contribution from non-synonymous coding variants to the changes in protein expression level, and the utility of these protein QTL variants in prioritizing GWAS risk genes for psychiatric disorders (*22*).

Another potential benefit of taking a multi-omics approach for identifying disease risk genes rests in the potential to dissect the fine details of regulatory mechanisms driving the disease-genotype association. Having relevant datasets to illuminate the origin and propagation of genetic impact could potentially arrive at a driver regulatory process from a risk variant to a risk gene. Ribo-seq is a technology that fills in the gap between transcript and protein expression. By adapting RNA-Seq to a ribosome footprinting method, ribo-seq provides transcriptome-wide quantification of ribosome occupancy (*25*, *26*), which can serve as a proxy for the amount of active translation synthesizing proteins from each mRNA transcript. When analyzed in conjunction with RNA-Seq and quantitative proteomics data, ribo-seq enables identification of translational and post-translational regulatory events (*20, 27*), both major steps defining the Central Dogma of molecular biology.

As a part of the consortium efforts to improve our understanding of the genetic basis of neuropsychiatric disorders (*28*), we generated multiple data modalities that included SNP genotyping, RNA-Seq, ribo-seq, and proteomics of postmortem cortical tissue samples of the BrainGVEX cohort, which altogether covered the major steps of the Central Dogma. In conjunction with the quantitative proteomics and transcriptome profiling results that we previously published (*22*, *29*), here we integrated ribo-seq data as our operational definition for protein synthesis, based on the ribosome occupancy level across the transcriptome, to advance our multi-omics investigation. Our results reveal regulatory properties of common variants and their utility in identifying the regulatory processes in the human brain driving disease risk for brain-associated complex disorders. It offers an opportunity to dissect and appreciate the regulatory information flow in the biological processes from a population perspective. Additional “rules” including an effect attenuation process is recognized.

## Results

### Measuring transcriptome-wide ribosome occupancy level in prefrontal cortex of adult human brain to quantify the level of protein synthesis

To investigate regulatory impact of genetic variants on protein synthesis in the prefrontal cortex of the human brain, we performed ribosome profiling on 211 prefrontal cortex samples from the BrainGVEX collection. In total we collected ∼62 billion ribosome profiling reads. Consistent with the expected ribosome footprint size, we found the average insert size of our ribo-seq libraries to range between 27.4 and 29.5 nucleotides. Similar to prior published studies (*30*), we found on average 74 % of unwanted reads from ribosomal RNA, tRNA, and snoRNA, which contributed no information to the translation of protein coding genes. After removing these unwanted reads, we obtained an average of 30.3 million uniquely mapped informative reads per sample (inter-quartile range: 20.5 ∼ 37.6 million reads) (Fig. S1A). Focusing on the informative reads, we found the majority (i.e. 84%) of our ribo-seq reads mapped to coding sequence (CDS) exons (Fig. S1B). Moreover, when visualized in aggregate across annotated coding genes, we found our ribo-seq data to show strong sub-codon periodicity at the expected positions (Fig. S1C). High proportion of CDS exon reads and strong sub-codon periodicity indicates the enrichment of footprints from ribosomes actively engaged in translating mature mRNA and reflects the quality of the dataset.

### Multi-omics cis-QTL mapping identified candidate regulatory variants and revealed translational and post-translational attenuation of eQTL effects

To identify variants associated with inter-individual expression differences, we perform cis-QTL mapping for each omics type independently. From the full dataset we filtered out low coverage genes and removed potentially low quality samples (see details in Data processing section in Materials and Methods) to reach a mapping dataset of 416 RNA-Seq samples, 195 ribo-seq samples, 268 proteomics samples, and the corresponding genotype data. Using the mapping dataset, we identified 12,411 eQTLs (out of 16,540 genes we deem sufficiently quantitated), 2,849 rQTLs (out of 14,572 genes), and 1,036 pQTLs (out of 8,013 genes) at FDR < 0.1. Overall, we observed similar genomic feature enrichment patterns of QTL variants across the three omics types (Fig. S2). On the other hand, when focusing on coding variants, we found pQTL to have the largest proportion of coding variants (pQTL: 11.5% vs. eQTL: 3.4% vs. rQTL: 5.1%). Among the coding variants, we found 82.4% were missense variants for pQTL comparing to 56.2% for rQTL and 46.4% for eQTL (Fig. 2A). Of note, we found three start loss variants (i.e. nonsynonymous substitutions in start codons) in rQTL, which was not found in eQTL.

**Fig. 1.**
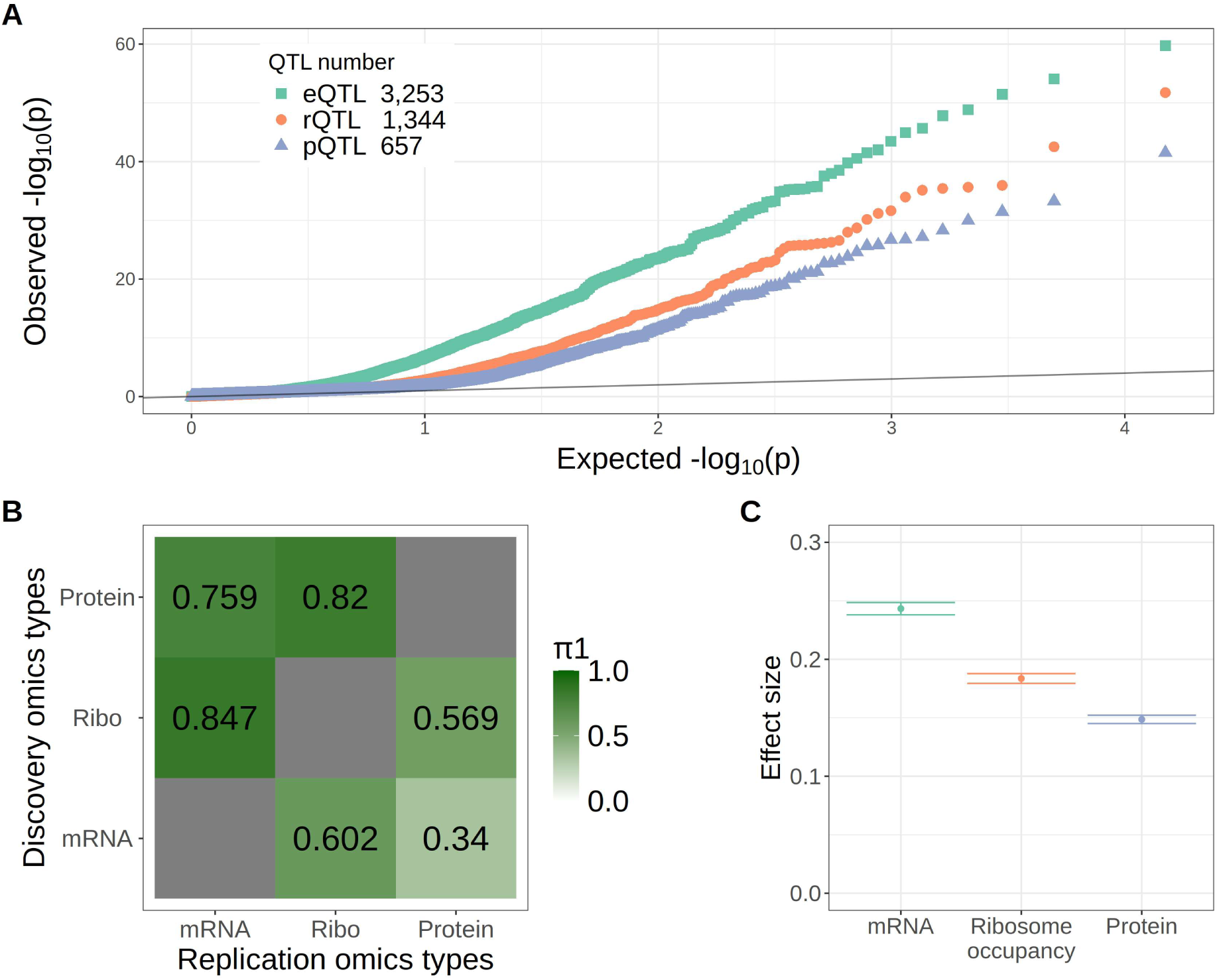
Genetic regulation of gene expression in the human brain. **(A)** P-value quantile-quantile plot between the observed (Y-axis) and the expected based on null distribution (X-axis). The black line indicates the expected distribution of p values when there are no real QTL signals. The number of cis-QTLs (i.e. the most significantly associated SNP for each gene) identified at 10% FDR is labeled in the top left inset. **(B)** Replication rate between QTL types. Proportions of QTLs replicated in the other two omics types are listed in the 3 × 3 matrix. Each row is a discovery omics type and each element of the row correspond to the proportion QTL signals replicated in the omics type specified by the column label. For example, only 34% of the eQTL signals were replicated in the protein data. Ribo: ribosome occupancy. **(C)** Effect size of CMC eQTL SNPs in BrainGVEX data. Mean and 95% confidence interval of absolute per allele effect across all CMC eQTL SNPs that were also analyzed in the BrainGVEX union set is shown.

**Fig. 2.**
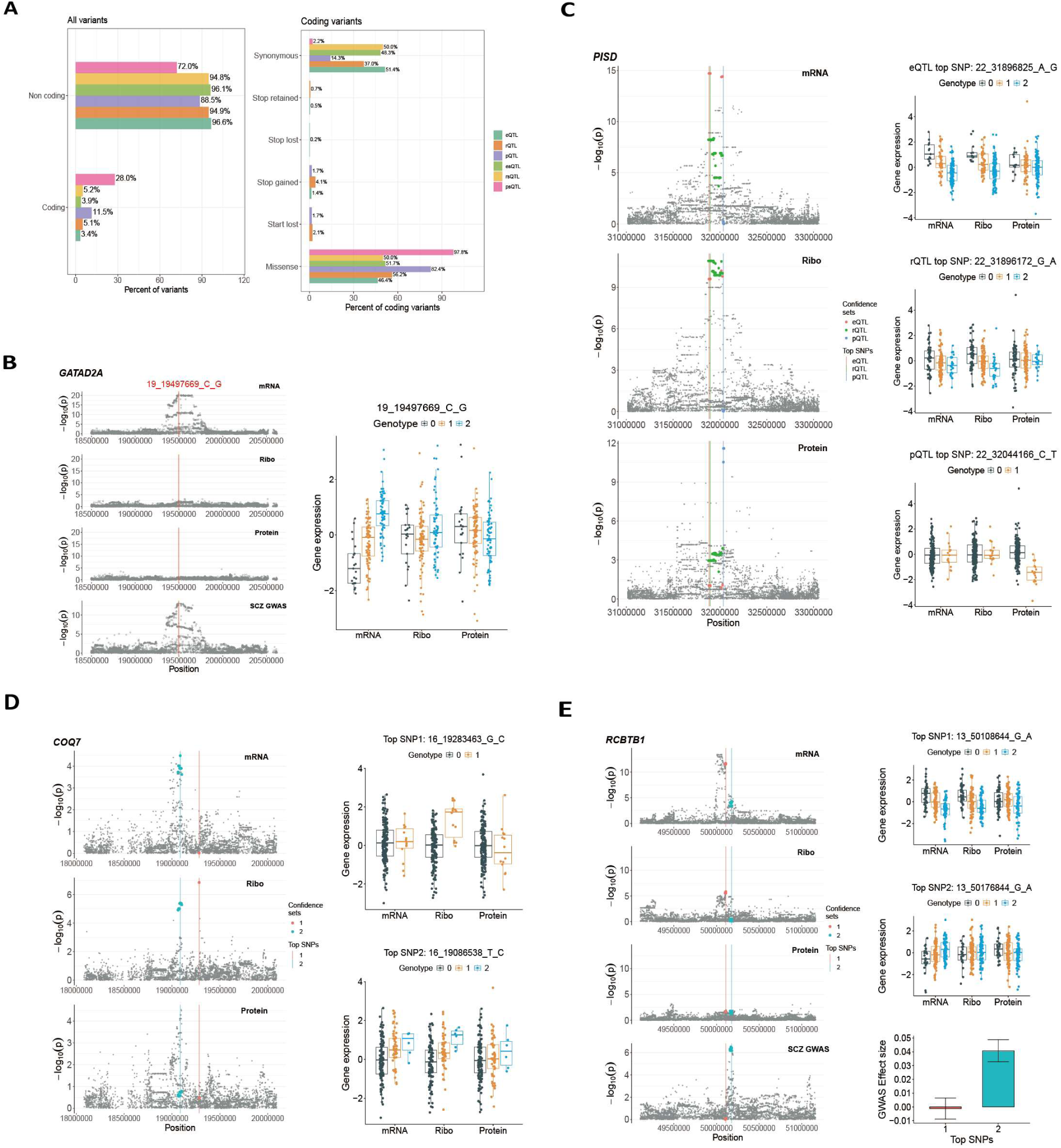
Omics-specific genetic regulations. (**A**) Categorizing QTL SNPs according to genic feature annotations. Bar plots showing the percentage of QTL SNPs in each category out of the total number of QTL SNPs considered. Note the high percentage (28%) of coding variants in psQTL SNPs out of all psQTL SNPs and the dominating percentage (97.8%) of missense variants in psQTL SNPs out of coding psQTL SNPs. (**B**) An example SCZ risk gene, *GATAD2A,* with predominantly transcript specific QTL effects. The red line indicates the top signal colocalization SNP between eQTL and SCZ GWAS on a Manhattan plot. Boxplots to the right summarize the normalized gene expression level stratified by eQTL SNP genotypes. Ribo: ribosome occupancy. (**C, D, E**) Example genes containing multiple QTL signals of varying omics specificity. Individual confidence set SNPs containing 95% of the QTL signal are color coded on a Manhattan plot with a vertical line indicating the top SNP that contributes the most signal. Boxplots to the right summarize the normalized gene expression level stratified by genotypes of the top SNP. Note that the example gene in (E) is a SCZ risk gene with two confidence sets. Only one of the two confidence sets’ QTL signal colocalizes with SCZ GWAS signal. The bar plots underneath the expression level boxplots show the log odds ratio +/- standard errors for the association between the confidence set top SNP genotype and SCZ, which is calculated based on GWAS summary statistics from Trubetskoy et al.

Intriguingly, we found drastic differences between omics types in the numbers of QTLs mapped with far fewer rQTLs and pQTLs identified, suggesting reduction of genetic effect along the path of Central Dogma. However, the differences in the number of genes tested between omics types and the differences in sample size complicate the interpretation. To better compare the effects of genetic regulation across data modalities, we identified 185 samples with 7,458 genes that were sufficiently quantitated across all three omics types (Fig. S3). Using this subset of data, we found 3,253 eQTLs, 1,344 rQTLs, and 657 pQTLs at FDR < 0.1 (Fig. 1A, Fig. S4, Table S1, S2, S3). Consistently fewer significant QTLs were identified as we moved downstream the Central Dogma of molecular biology.

A challenge in comparing between the numbers of QTLs identified from each omics type rests in the fact that not all true effects were identified. While the majority of the QTLs identified here were replicated in independent datasets (Table S4, S5; Fig. S5), which provides an experimental support for our discoveries, the level of false negative rates is less clear. Tests replicating QTLs identified from one omics type in the other omics types can better capture the proportion of genetic effects shared between QTL types. We performed replication tests using π1 estimates from the qvalue method (*31*). Overall, we found high proportion of QTLs replicated in other omics types (Fig. 1B). However, when considering the replication rates with the direction of genetic information flow, we found asymmetric replication rates, with the downstream omics types to replicate less than the upstream omics types. More specifically, we found 84.7% of the rQTLs were replicated at the transcript level, but only 60.2% of the eQTLs were replicated at the ribosome occupancy level. Moreover, while 75.9% of the pQTLs were replicated at the transcript level, only 34.0% of the eQTLs were replicated at the protein level (Fig. 1B). A similar asymmetry in proportion replicated between upstream and downstream omics types was observed when using a direction-aware cutoff-based approach across a wide range of significance cutoffs (Fig. S6). The lower percentages of eQTLs and rQTLs replicated at the protein level indicate potential effect attenuation (i.e. either the inter-individual variation in gene expression becoming smaller and therefore harder to detect or a lack of such effect in the downstream omics types). Such effect attenuation at the protein level has been previously reported in lymphoblastoid cell lines (LCLs) (*20*). Our brain data revealed an additional layer of complexity, where effect attenuation was also observed at the level of mRNA translation.

While our replication tests revealed a trend of effect attenuation for eQTL variants in the downstream phenotypes (Fig. 1B), these same observations could alternatively be explained by differences in statistical power between technologies. An independent piece of evidence that is not sensitive to measurement precision is needed to reach a solid conclusion. Fold change estimates from less precise measurements are noisier but not biased. In other words, by directly comparing QTL impact on expression level change across omics types we could gain a direct view of effect size attenuation. One issue that remains is that using eQTLs identified from our dataset could bias the results by artificially inflating the variant pool with large eQTL effect sizes, which might arise simply from applying the QTL mapping procedure to the same dataset (i.e., a technical ascertainment bias). Using eQTLs independently identified from prefrontal cortex samples by the CommonMind Consortium (CMC) (*32*), we avoid the ascertainment bias and can therefore compare the effects size of eQTL variants between the three omics types using our dataset. A similar approach was successfully implemented to address this power issue in previous work in LCLs (*20*). Using 5,915 CMC eQTLs that were also quantitated in our dataset, we found the eQTL effects on the transcript expression level to be significantly larger than their effects on ribosome occupancy level (per allele log_2_ fold differences: mRNA 0.2433 [95% CI= 0.2381∼0.2486] vs. ribosome occupancy 0.1836 [95% CI= 0.1794∼0.1878]), which were in turn significantly larger than their effects on protein level (per allele log_2_ fold differences: 0.1486 [95% CI= 0.1451∼0.1522]) (Fig. 1C, t-test P < 2.2e^-16^ for all pairwise comparisons). Moreover, for these CMC eQTLs, we found translational regulation to account for more effect size reduction than post-translational regulation (Fig. 1C). By focusing on the effect sizes of independently identified eQTLs, our results strongly support the presence of downstream mechanisms attenuating eQTL effects both at the ribosome occupancy level (translationally) and at the protein level (post-translationally). A complementary analysis looking at the effect size of pQTL variants identified from an independent study (*33*) further supported our conclusion. We found, in our dataset, larger effect size at transcript level when comparing to the fold change at protein level for these independently identified pQTL SNPs (Fig. S7). We noticed that for these independently identified pQTL SNPs, the eQTL effect attenuation appears to be predominantly translationally regulated (Fig. S7B; mRNA vs. ribosome occupancy t-test P = 6.6e^-15^ and ribosome occupancy vs. protein t-test P = 0.4). Similar results were observed across a wide range of significance levels used to define pQTL (Fig. S7).

In contrast to prior studies in LCL (*20*), we found clear translational attenuation of eQTL effects in prefrontal cortex. To further investigate this apparent discrepancy, we cross referenced the genes associated with eQTL identified in prefrontal cortex samples to those identified in LCL. We found rather limited overlaps: of the 3,253 eQTL associated genes identified in our brain data only 598 (i.e. ∼18%) were found associated with a significant eQTL in LCL (Fig. S8A), indicating the possibility that the between study differences in eQTL discoveries could play a role in the observed differences in translational attenuation of eQTL effects. Indeed, when inspecting the downstream effect of LCL specific eQTL versus a set of effect size matched brain specific eQTL in our brain dataset we found no attenuation of eQTL effect at the ribosome occupancy level among LCL specific eQTL while translational attenuation of eQTL effect remains significant among brain specific eQTLs (Fig. S8B). On the other hand, the shared eQTL found between the two studies showed the expected translational attenuation of eQTL effects in the brain dataset (Fig. S8C), again supporting the notion that the distinct groups of eQTL analyzed between the two studies drive the observed discrepancy.

### Identifying omics specific QTLs and their signal colocalization with brain disorder GWAS

The prevalent effect size reduction of eQTLs raised the question of the relevance of these genetic regulation at the organismal level. Because most cellular tasks are executed by proteins, the genetic regulatory effects not reaching the protein level are, presumably, less likely to have an impact on organismal traits. More specifically, we seek the biological relevance of expression specific and ribosome occupancy level specific QTLs (i.e., esQTL and rsQTL). To answer this question, we first set out to investigate the relevance of different QTL types in brain associated complex traits such as neuropsychiatric and neurodevelopmental disorders. By applying mediated expression score regression (MESC) (*34*) to summary statistics from GWAS of four different complex brain disorders and our three types of molecular QTL summary statistics we estimated for each disorder the proportion of heritability mediated by each of our molecular QTL types (Table S6 and Fig. S9). We noticed that for schizophrenia (SCZ), bipolar disorder (BD), and autism spectrum disorder (ASD), eQTL mediated the most heritability, followed by rQTL, which is then followed by pQTL. For example, using the Trubetskoy et al. SCZ GWAS (*18*), we found our eQTLs to mediate 7.09%, rQTLs to mediate 4.06%, and pQTLs to mediate 2.17% of SCZ heritability (Table S6). On the other hand, for major depressive disorder (MDD) all three QTL types mediated similar proportion of heritability (∼2.5%).

After establishing the relevance for each of the three QTL types in brain associated disorders, we next sought to identify omics-specific QTLs, in order to further evaluate their relevance in driving disease risk. For example, we aim to identify expression specific QTLs (i.e. genetic variants that impact transcript level of the linked genes but not the downstream ribosome occupancy level nor protein level) that colocalize with GWAS signals of brain associated disorders. To identify omics-specific QTLs we took a multiple regression approach by first removing effects of the two other omics-types and then test for genotype association with the residual. To further reduce the number of false positives, for each potential omics-specific QTL identified at 10% FDR we further test genotype association with the other two omics-types and remove any variant that has nominally significant association. We identified 1,553 loci/genes with expression specific QTL (esQTL), 155 genes with ribosome occupancy specific QTL (rsQTL), and 161 genes with protein specific QTL (psQTL) (Fig. S10). Similar to standard QTLs we found enrichment of esQTL and psQTL in genomic features such as the exons and the 3’UTRs (Fig. S2). In addition, we found even higher proportion of coding variants and missense variants among psQTLs comparing to the enrichment of these categories reported above for pQTL (Fig. 2A). Of note, we found a clear downward shift in 5’UTR enrichment from pQTL to psQTL (odds ratio 2.51 vs. 1.52, P < 5e^-6^). Similar decrease in 5’UTR variant enrichment was also observed among the other two pairs of standard and omics-specific QTL (i.e., eQTL vs. esQTL and rQTL vs. rsQTL) indicating the possibility that QTL variants in 5’UTR are more likely to be shared across omics types, presumably because they impact protein expression level via transcript level and/or ribosome occupancy level.

To identify genes with omics-specific QTL effects driving disease risk, we perform, for each gene containing omics-specific QTL signal, *coloc* analyses with default prior (*11*) between the nominal p value from *cis-*QTL mapping and each brain associated disorder GWAS. We performed separate analysis using either the nominal p value from the omics-specific *cis-*QTL mapping or the standard *cis-*QTL mapping. At a colocalization posterior probability cutoff of 70% we identified risk genes for each omics-type by brain disorder combination for both approaches (Table S7). We found the most risk genes from esQTL loci, with 14 genes showing signal colocalization between SCZ GWAS signals and either standard or omics-specific eQTL signals, and 14 genes for BD, while the remaining combinations resulted in single digit colocalizations (Table S7). For the majority of the risk genes, we observed consistent results between the two approaches (Table S7). On the other hand, we found a few risk genes that have GWAS colocalization with only standard QTL signals, indicating the possibility of GWAS signals colocalizing with QTL signals that were shared with, and therefore regressed out by, other omics types at these loci. In other words, some of these omics-specific QTL were identified from loci containing shared QTL signals that colocalized with GWAS signal.

Because regulation of a gene is often modulated by multiple genetic variants, to evaluate the consequence of overall *cis*-QTL impact on gene expression, we use PrediXcan to estimate aggregate genetic regulatory effects for each gene. To identify genes that have omics-specific QTL effects, for each omics type we built PrediXcan models using normalized quantifications with and without regressing out signals from the other omics types. A significant PrediXcan model fit on residuals indicates the potential presence of omics-specific QTL signals (i.e., QTL signals remained after regression). Using a high PrediXcan model fit R^2^ cutoff of 0.1 we identified 1,098 candidate genes containing omics-specific QTL signals. To further identify genes with predominantly omics-specific QTL effects we then remove, from the candidate gene list, genes that have either shared QTL effects or additional QTL effects from other omics types by filtering out genes with significant PrediXcan model built from any of the two other omics-types. Using this approach we identified 382 esQTL genes, 36 rsQTL genes, and 51 psQTL genes that have predominantly omics-specific QTL effect. Consistently, these predominantly omics-specific QTL effect genes identified using the PrediXcan gene-based approach are enriched of omics-specific QTL variants identified from the SNP based multiple regression approach (Fig. S11). Colocalization with brain-associated complex disorder GWAS identified 11 risk genes with predominantly omics-specific QTL effects (Table S8). We found most signal colocalization with SCZ, which resulted in 7 SCZ risk genes with predominantly esQTL effects (Fig. 2B).

For the remaining 629 candidate genes that potentially contain both omics-specific and shared QTL signals or contain independent omics-specific signals from multiple omics types (i.e. the candidate omics-specific QTL effect genes filtered out from the above step), we use SuSiE to identify independent QTL signals from each omics for each gene (i.e. SuSiE 95% confidence sets). By separating out independent signals form the same gene using SuSiE we can then operate on the confidence set level to identify additional omics-specific QTL signals that colocalize with complex disorder GWAS. We identified a total of 829 confidence sets from 594 candidate genes (Fig. S12). For each confidence set, we perform QTL signal colocalization between omics-types using *coloc*. We found 404 confidence sets to contain QTL signals that are shared with at least one other omics types (i.e. PPH4 > 0.7) while 96 contain QTL signals that are shared across all three omics types (Table S9 full table). Of note, of the 594 candidate genes we found 170 genes to have at least two confidence sets. Among these genes with multiple confidence sets we found examples of separate omics-specific QTL signals from different omics types (Fig. 2C) and example genes that contain both omics-specific and -shared QTL signals (Fig. 2D).

To further investigate the relevance of these QTL confidence sets in brain-associated disorders we search for colocalization with brain-associated disorder GWAS signals. Of the 829 confidence sets we found 25 (from 25 genes) to have signal colocalization with at least one of the four brain-associated complex disorders tested. Of these 25 GWAS colocalization signals 13 have QTL signals shared between at least two omics types. Upon visually inspecting the nominal p values from QTL mapping and brain associated disorder GWAS signals at these risk loci we found a few genes with apparently differential colocalization results between confidence sets. For example, we found *RCBTB1* to have two eQTL confidence sets while only one of the two confidence sets contain eQTL signals colocalizing with SCZ GWAS (PPH4 = 0.8). Interestingly, the GWAS colocalizing eQTL for the *RCBTB1* locus appears to be omics specific while the other confidence set appears to contain a QTL signal that is shared between transcript expression and ribosome occupancy (PPH4 = 0.7) (Fig. 2E).

### Functional genomics identification of Schizophrenia risk genes

To further investigate the relevance of attenuated eQTLs in brain disease risk, we next took a complementary approach by first identifying risk genes from each omics type separately and then investigating the relevant regulatory processes driving disease risks. Given that we observed the highest number of risk genes for SCZ from our omics-specific QTL analyses, here we focused on SCZ. We applied S-PrediXcan (*35*) on the Trubetskoy et al SCZ GWAS data and our subset of multi-omics data. At 5% family-wise error rate, we found 52, 29, and 16 SCZ risk genes, respectively from RNA-Seq, ribo-seq, and proteomics data (Fig. 3A, Fig. S13). Consistently, when comparing the corresponding GWAS-QTL *coloc* results between genes that passed and genes that failed the S-PrediXcan tests, we found S-PrediXcan risk genes to have higher colocalization posterior probability (Fig. S14). Among S-PrediXcan risk genes only four genes, *NEK4*, *KHK*, *CNNM2* and *DARS2* were consistently identified as SCZ risk genes from all three omics types (Fig. 3B). The majority (74.3%) of the risk genes were identified only from one of the three omics types. This limited sharing in risk gene identification between omics types is consistently observed across multiple significance levels and is in clear contrast to the amount of signal sharing found between QTL types (Fig. 1B; Fig. S15; Table S11, S12, S13).

**Fig. 3.**
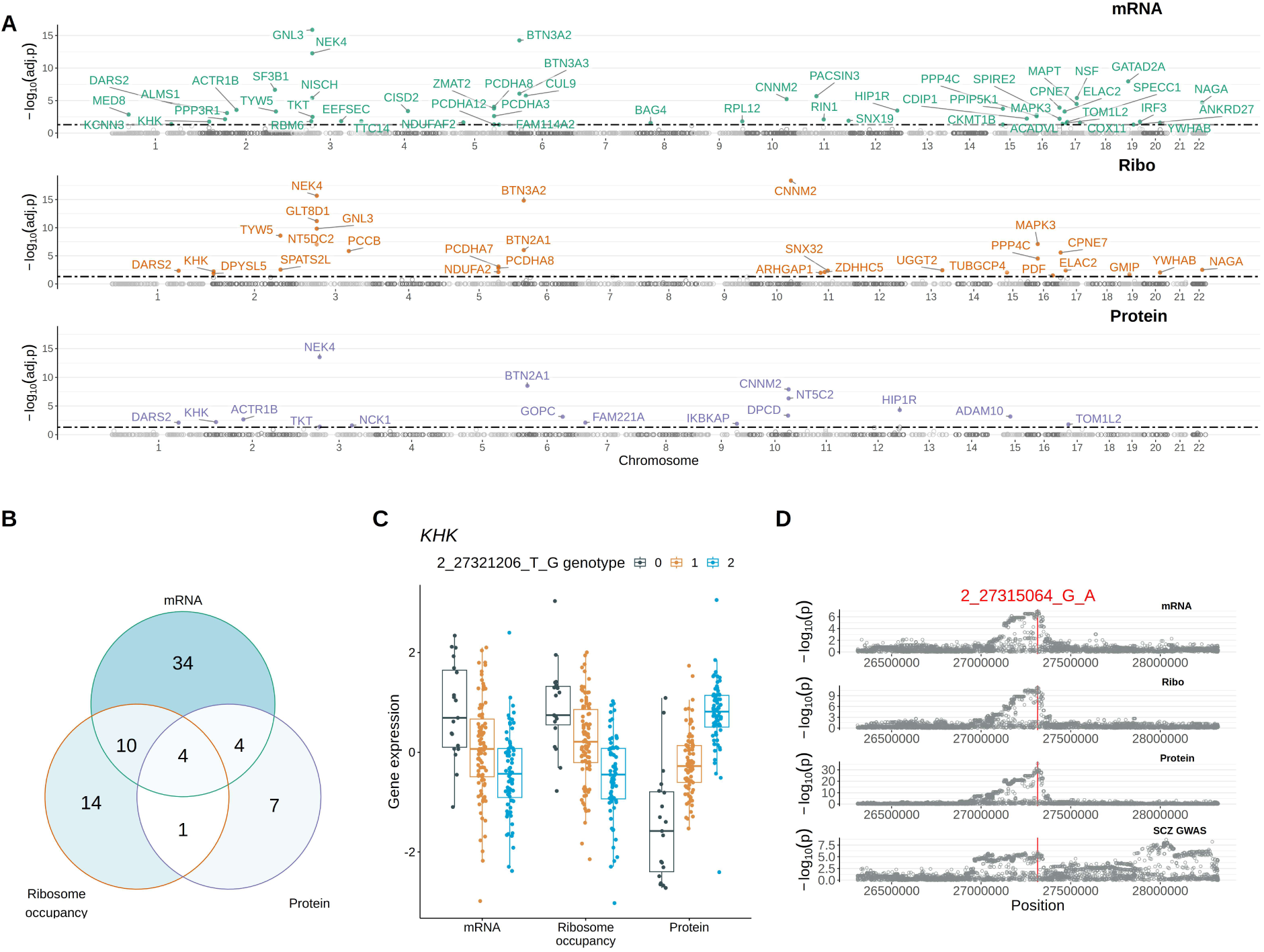
Schizophrenia risk genes identified from each of the three omics types, RNA-Seq (mRNA), ribo-seq (ribosome occupancy), and quantitative mass spectrometry (protein), using S-PrediXcan. **(A)** Manhattan plots showing significance level (i.e. -log10 FWER from S-PrediXcan) of gene expression-schizophrenia association across the genome for genes that pass the 5% FWER significance cutoff for each omics type. The black horizontal dotted line indicated the significance cutoff (5% Bonferroni-adjusted p value). Ribo: ribosome occupancy. **(B)** Venn diagram illustrating the number of overlapping risk genes identified between omics types. **(C)** Boxplots summarizing normalized gene expression level stratified by eQTL SNP genotypes for *KHK*. **(D)** Manhattan plots showing nominal p value distribution for each QTL type and schizophrenia GWAS for the 1Mb QTL mapping window flanking *KHK*. The red line indicates the position of the lead colocalization SNP between eQTLs and schizophrenia GWAS.

Among the 74 risk genes we identified using S-PrediXcan (i.e. the union of the risk genes identified from each of the three omics types), 44 have previously been reported in GWAS studies as either the mapped genes or as one of the nearby genes under the GWAS peak (Table S14) (*18*, *23, 29*, *36–39*). Of these previously reported genes, 27 matched the mapped genes while the remaining 17 nominated an alternative candidate gene for each GWAS locus (Note that for some of these 17 loci, the original GWAS signals were mapped to an intergenic region). Comparing our results to other published multi-omics SCZ risk gene identification studies, we found 15 matched to the risk genes identified by Giambartolomei et al., which used RNA-Seq and DNA methylation data from prefrontal cortex of the human brain (*23*), and 12 matched to the 120 prioritized SCZ risk genes from Trubetskoy et al., which used colocalization with eQTL and Hi-C data (*18*). When comparing our results to other published TWAS SCZ risk gene identification studies, we found 19 matched to the risk genes identified by Gandal et al. (*29*), and 21 matched to the risk genes identified by Gusev et al. (*39*) (See Supplemental Table S14 for an extended list of risk gene comparisons to prior TWAS studies). On the other hand, for 22 of our 74 risk genes (30%), we found no match to the known risk gene list (Table S14, bottom rows), which we compiled based on previous GWAS, TWAS, and other functional genomics risk gene identification studies (*18*, *23*, *29*, *36–39*). These “no match” novel risk genes have relatively weak SCZ GWAS signals and are therefore challenging to identify without the additional information provided by our multi-omics QTL dataset. For example, we found strong colocalization between a modest SCZ GWAS signal at *2p23.3* and all three types of molecular QTLs of the gene *KHK* (Fig. 3D). Intriguingly, the *KHK* pQTL is in opposite direction of the *KHK* eQTL and rQTL (Fig. 3C), suggesting a strong post-translational process regulating its protein level in the opposite direction to the transcriptional regulation. *KHK*, Ketohexokinase, plays a pivotal role in fructose metabolism and has been hypothesized to contribute to neuronal glycolysis and the loss of neuronal functions (*40*). In addition to *KHK*, other novel SCZ risk genes have also been reported to participate in neuronal/ cognitive functions: For example, in neuroglial cell lines, silencing of *SNX32* leads to defects in neuronal differentiation (*41*). As another example, *NSF*, N-Ethylmaleimide Sensitive Factor, which encodes a vesicle fusing ATPase, was reported as a causal factor in intelligence traits (*42*).

### Analyses of multi-omics dataset reveal regulatory mechanisms of schizophrenia risk genes

While PrediXcan is a powerful tool for GWAS risk gene identification, it does not control for potential horizontal pleiotropy (*43*). To this end, we performed two-sample Mendelian Randomization (MR) with Egger regression to replicate the risk genes we identified from using S-PrediXcan. Egger regression includes an intercept term, which can be used to evaluate the level of horizontal pleiotropy (*44*). For each risk gene we first used LD-clumping (*45*) to identify the top QTL SNP from each clump (i.e. a group of linked variants defined by the LD cutoff) to serve as strong genetic instruments (*46*). We then tested for the causal relationship between gene regulation (i.e. QTL signal) and SCZ (i.e. GWAS). We used an operational definition of a causal effect based on the MR test results (see Methods). At 10% FDR, of the 97 gene-by-omics combinations (i.e. a total of 74 risk genes including some discovered from more than one omics type), 67.0% (65/97) passed the MR test. Of those 33.0% that failed the MR test, 37.5% (12 out of 32) failed because of horizontal pleiotropy identified by the Egger intercept test (Table S15). A total of 53 risk genes were replicated in at least one of the three omics types. Note that in order to increase our power, we used a relaxed R^2^ cutoff of 0.5 for LD-clumping and adjusted for the linkage between instrument variables in our MR tests using generalized IVW and Egger (*47*). Similar causal effects were observed when using either a stringent R^2^ cutoff of 0.01 or using SuSiE (the Sum of Single Effects) (*48*) as an alternative approach to select instrument SNPs (Fig. S16). However, these more stringent criteria on instrument variable independence led to lower power to detect and fewer genes available to test (Table S16). For example, when using SuSiE for selecting instrument variables, only 40 genes were tested because some of the risk genes had no fine mapped QTL SNPs according to SuSiE.

A key strength of using a multi-omics QTL approach to identify GWAS risk genes rests in the possibility of further narrowing down the potential regulatory mechanisms. To this end, we further examined the likely causal mechanisms for the 53 replicated SCZ risk genes using one-sample Mendelian Randomization to infer causality between QTL types. We focused our analysis on independently testing for causal effects between neighboring QTL types following the direction of information flow of the Central Dogma (i.e. mRNA -> ribosome occupancy, and ribosome occupancy -> protein). Here we used fine-mapped QTL SNPs identified from the exposure omics types (i.e. the upstream omics types) as instrument variables for one-sample MR analysis. Among the 53 MR-Egger replicated SCZ risk genes, we found 21 genes with significant causal effects both from eQTL to rQTL (i.e. the upstream pathway) and from rQTL to pQTL (i.e. the downstream pathway) (both-passed risk genes; Fig. 4A). Significant causal effects detected from both pathways suggest transcriptionally regulated protein level differences as the potential mechanisms driving these risk genes in SCZ etiology (see an example in Fig. 4B). On the other hand, 25 and 7 replicated risk genes have either significant causal effects only in one of the two pathways (single-passed risk genes; Fig. 4A) or have no significant causal effects (none-passed risk genes; Fig. 4A), respectively.

**Fig. 4.**
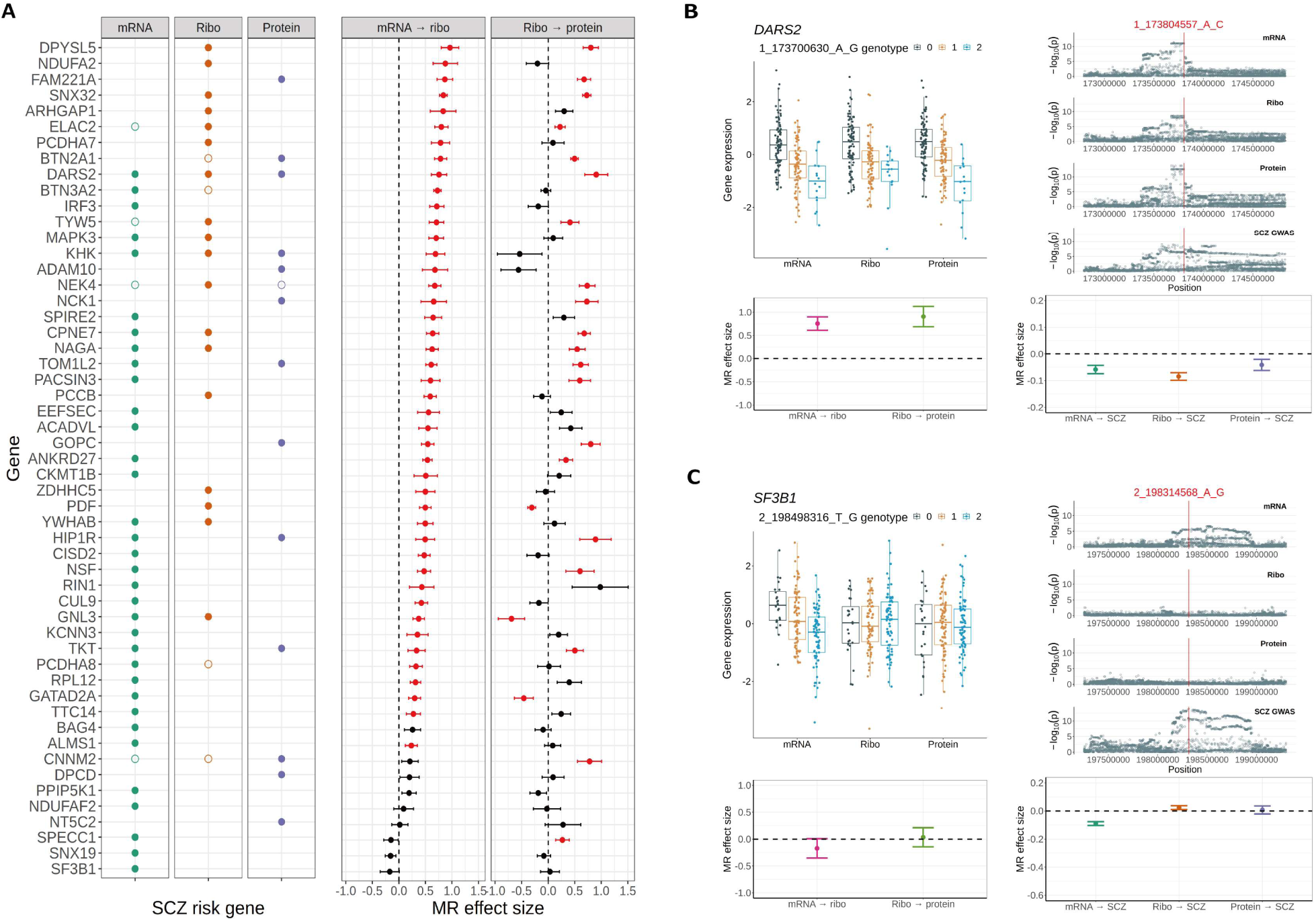
Identifying driver regulatory mechanism for SCZ risk genes using Mendelian Randomization (MR). **(A)** Summaries of MR test results. Left panel: Two-sample MR replication of S-PrediXcan SCZ risk gene discoveries. Each circle represents a S-PrediXcan discovery. A solid-colored circle indicates successful replication in a MR test. Right panel: One-sample MR test results for causality between omics types. Shown are MR effect size +/- the corresponding standard errors. Red data points indicate significant test results. Ribo: ribosome occupancy. (**B, C**) Example SCZ risk genes driven by either transcriptional regulation of protein expression (B) or transcript specific effects (C). Boxplots on the left summarize the normalized gene expression level stratified by QTL SNP genotypes. The effect size estimates from the corresponding one-sample MR causality test between each pair of omics are shown in the plot beneath. Manhattan plots on the right show nominal p values for all variants in the QTL mapping window for each omics and SCZ GWAS with the red vertical line indicating the top signal colocalization SNP between QTLs and GWAS. Directly underneath the Manhattan plots are effect size estimates from the corresponding two-sample MR tests replicating the risk gene discoveries.

Of note, for 23 of the 25 single-passed risk genes, we found significant causal effects in the upstream pathway (mRNA -> ribosome occupancy). This asymmetry is reminiscent of the eQTL effect attenuation described in the prior sections. A failed test could indicate either a true lack of effect or a lack of statistical power. To take a closer look, we directly assessed the effect size, the noise level, and instrument strength of the one-sample MR test results. When comparing to the 21 both-passed risk genes, we found clearly smaller effect sizes for the failed tests of both the 7 none-passed risk genes (average 0.577 vs 0.058, t-test P < 10^-3^, Fig. S17A) and the 23 single-passed risk genes (average 0.524 vs 0.116, t-test P < 10^-3^, Fig. S18A). Note that in both cases the inter-quartile range of the estimated causal effects for the failed tests covered zero (Fig. S17A, Fig. S18A). For these same comparison groups, we found no significant differences in instrument strength (Fig. S17C and S18C) and found slightly higher noise level in the failed MR tests (Fig. S17B and S18B). When considered together, these observations (i.e. clearly smaller effect size and slightly elevated noise level for the failed tests) indicate that many of these test results reflect a true lack of causal effect while in some cases weak effects were obscured by noise. In other words, many of these risk genes are likely to be driven by either omics-specific QTLs or attenuated eQTLs. Case in point, we found *SF3B1* to have strong signal colocalization between eQTLs and SCZ GWAS without clear QTL signals in either ribo-seq or proteomics data (Fig. 4C).

## Discussion

Using a panel of postmortem prefrontal cortex samples, we found clear evidence of post-transcriptional attenuation of eQTL effects in the human brain. Many of the differences found in transcriptomes were not present in proteomes. This observation echoes earlier work in HapMap lymphoblastoid cell lines (*20*) and extends the prior conclusion from *in vitro* cell lines to complex human tissues. Importantly, distinct from the earlier work in lymphoblastoid cell lines, which found rQTL to mostly track with eQTL, we found clear attenuation of eQTL effect in ribo-seq data indicating that translational regulatory processes are involved in eQTL effect attenuation in the human brain. Our further investigation on this observation indicated that differences in the set of eQTL (eGENE) identified between the two studies explains the discrepancy. However, it remains to be determined whether the discrepancy observed between the two studies reflects a true difference between the tissue/cell types used in these studies, or effects rooted in differences in sample complexity, or effects from other unknown unaccounted for variations, or simply an incidental finding. Prevalent translational attenuation of variant impact on transcriptional gene expression level has previously been reported in budding yeast (*49*, *50*). However, results from follow up studies (*51*) appear to present an inconsistent picture. Here, using replication tests for individual eQTLs and testing for aggregate effect size of eQTLs independently identified from CMC, we provided strong evidences supporting prevalent translational attenuation of eQTL effects in the human brain. Importantly, by focusing on comparing QTL effects (i.e. between genotype fold change) in aggregate, our analyses are robust to power issues resulted from measurement imprecision (random noise). On the other hand, our conclusions could still be sensitive to potential biases in the dataset, which could result from technical issues that cannot be randomized. For this reason we paid close attention to the impact of sequencing coverage (for sequencing data) and ratio compression (for quantitative mass spectrometry data) on our major conclusion. Of note, our quantitative mass spectrometry study employed an analytical procedure that adjusts for contaminating background signals in order to correct for potential ratio compression in quantification (*52*). Consistently, among low abundance proteins, which are more susceptible to the influence of ratio compression, we did not find a disproportional decrease in pQTL effect size (Fig. S19C). Similarly, while genes with low expression level are more sensitive to issues related to low sequencing coverage, we found little to no correlation between eQTL/rQTL effect size and expression level (Fig. S19A, S19B). Although ample precautions and multiple distinct approaches were taken to confirm the observed effect size attenuation, it remains possible that unknown, unaccounted for omics-specific features could potentially confound the results. In other words, although operating at the molecular level, our study remained observational. Future work replicating this observation and elucidating molecular mechanisms of post-transcriptional attenuation of eQTL effects are needed to provide a clear understanding of the phenomenon.

Although our study was not designed to answer the key questions regarding the underlying mechanisms of the observed translational and post-translational attenuation of eQTL effect, the high prevalence of the observed eQTL effect attenuation when considered in conjunction with the low prevalence of coding mutations in human population provides some hints. Since a compensatory variant driving translational or post-translational attenuation is expected to be either a coding mutation that impact translation elongation rates or protein turnover rates or a UTR variant that impacts translation initiation rate, the apparent mismatch in incident prevalence indicated that compensatory variants are unlikely to explain the majority of the effect size attenuation and therefore implies a broad-spectrum mechanism that reduce variation at the protein level across genes. Consistently, we observed significant translational and post-translational attenuation of eQTL effect across all PhastCons score strata on top of a clear trend of decreased QTL effect size as the evolution conservation scores increase (Fig. S20). Our current work focuses on the other important follow-up question of whether the attenuated eQTLs are functionally (biologically) relevant. We attempted to address this question by exploring the relevance of attenuated eQTLs in brain associated disorders, such as SCZ, a neuropsychiatric disorder that is highly heritable. We took multiple approaches to identify risk genes that have either omics-specific QTL signals or attenuated eQTLs. In one approach, we used regression based methods to identify omics-specific QTL performing separate analysis either at the SNP level or using PrediXcan to aggregate variant impact at the gene level. In both cases, we found the most number of colocalization between esQTL and complex disorders (i.e. comparing to rsQTL and psQTL). From the gene level analysis we further identified interesting example genes containing multiple QTL signals with varying level of omics-specificity, including an example gene, *RCBTB1.* We found two strong eQTL signals at *RCBTB1*, while one potentially shared and one potentially omics-specific. Only the apparently omics-specific eQTL was found to colocalize with SCZ GWAS. These examples illustrate the complexity of variant impact on gene regulation and disease risks. Regression based approach, however, could produce false positive omics-specific QTL simply from noisy predictors. Although we implemented checks and filters to remove these likely false positives, this caveat is important to keep in mind when interpreting these results.

In another approach, we conducted our search by first identified risk genes from all quantitated genes using a TWAS approach, S-PrediXcan method. We then tested for causal regulatory mechanisms driving the risk gene using QTL SNPs as instrumental variable in a one-sample Mendelian Randomization test. Before testing for causal regulatory mechanisms, we first replicated the TWAS risk genes using two-sample Mendelian Randomization with Egger intercept tests. About 30% of the risk genes that we identified here have not previously been reported as SCZ risk genes. The use of our functional genomics dataset is key to the discovery of these novel SCZ risk genes because of their relatively weak GWAS signals (see Fig. 3D for an example). As a reminder, we would like to emphasize the fact that individual risk gene identification is dependent on the choice of significance cutoffs, which is to some extend arbitrary. We used a rather stringent family-wised error rate for risk gene identification in order to reduce the number of false positive results that could mislead researchers interested in molecular biology follow-up studies.

To identify causal regulatory mechanisms for each replicated risk gene, we further tested causality between QTL types using one-sample MR. We found 21 risk genes that are likely contributing to SCZ risk through transcriptionally regulated protein level. On the other hand, we also identify 7 genes that show no significant results from MR tests (candidate omics-specific risk genes) and 23 genes that have significant MR test results only from transcription to translation (candidate post-translationally-attenuated risk genes). In essence, attempts to identify omics-specific risk genes (or omics-specific QTLs) are dealing with the challenge of separating true negatives from false negatives. As such, the interpretation of the failed MR tests is challenging. Our subsequent analyses looking at comparing instrument strength, noise level, and effect size between passed and failed MR tests, indicated comparable instrument strength, slightly elevated noise level in the failed tests, and clearly smaller effect size in the failed tests. In other words, it remains possible that some of the failed tests could be reflecting a small effect size obscured by the elevated noise level. In addition to the issues with false negative results, some of the discoveries could be false positives to begin with. Although we have good confidence with our FDR and FWER estimates, given that our test statistics for QTL mapping and risk gene identification are well calibrated (Fig. S4, S13), pleiotropy could introduce positive results from a different underlying cause (i.e. true effects on SCZ risk but false positive risk genes).

Power issues, however, do not explain the whole story. As was consistently observed throughout our study, when viewed in aggregate, we see clear effect size differences between omics types, both for QTLs and for causal effects from MR tests. These effect size estimates are not influenced by significance cutoffs and are not biased by power differences. Such general trends are also unlikely to simply be a result of spurious associations or made up entirely of false positives. Our results therefore bring forth an interesting mechanistic question: how do attenuated eQTL variants impact SCZ without changing protein levels? One interesting possibility is that the biologically relevant traits here are protein synthesis rates and protein turnover rates. An association between genotype and protein synthesis rates could manifest in the form of an attenuated eQTL, if the differences in per transcript protein synthesis rate (e.g. differences in ribosome elongation rate or translation initiation rate) appears to offset the differences introduced at the transcript level, which in turn could resulted in a lack of differences in ribosome occupancy level. Similarly a protein turnover QTL could also manifest in the form of a rQTL attenuated at the protein level. In other words, colocalization between an attenuated rQTL and a GWAS signal could actually be reflecting a colocalization of the GWAS signal with a protein turnover QTL. Testing this hypothesis requires a direct measurement of protein turnover rate. We hope by presenting the current results, our findings can inspire future studies on this topic to better understand the regulatory processes from DNA to brain associated disorders.

## Materials and Methods

### Data sources

The SNP genotypes (*22*), RNA-Seq (*29*), and quantitative mass spectrometry (*22*) data generated for prefrontal cortex tissue samples of the BrainGVEX cohort were downloaded from the PsychENCODE Synapse portal (https://www.synapse.org/#!Synapse:syn5613798) (See Table S18 for a summary on the number of samples in each dataset and their respective overlap with the samples in the genotype data). The BrainGVEX cohort includes 420 Caucasians, 2 Hispanics, 1 African American, 3 Asian Americans, and 14 unknown-origin individuals. We also used GWAS summary statistics data of schizophrenia (*18*), bipolar disorder (*53*), autism spectrum disorder (*54*), and major depression disorder (*55*) obtained from the Psychiatric Genomics Consortium.

### Ribosome profiling

Ribosome Profiling experiments were performed using Illumina’s TrueSeq Ribo Profile (Mammalian) Kit. TrueSeq Ribo Profile (Mammalian) Kit was developed for cell lines. We adapted the protocol to frozen tissue samples with MP Biomedicals™ FastPrep-24™ Classic Bead Beating Grinder and Lysis System. Specifically, 60-80 mg of frozen brain tissue was homogenized in Lysing Matrix D tubes containing 800 µl polysome lysis buffer. The Lysing tubes were placed on the FastPrep-24™ homogenizer set at 4.0 m/s with 20 s increments until no visible chunks of tissue remained. Tissue lysate was incubated on ice for 10 min followed by centrifugation at 20,000g at 4 °C for 10 minutes to pellet cell debris. Clarified lysate was subsequently used for ribo-seq library preparation following TrueSeq Ribo Profile (Mammalian) Kit instructions. Indexed libraries were pooled and then sequenced on an Illumina HiSeq 4000. Note that the experimental protocol for TrueSeq Ribo Profile (Mammalian) Kit that we followed is a modified version of the previous ribo-seq protocol published by Ingolia and colleagues (*26*), and it has the following key modifications: Monosome isolation was performed using Sephacryl S400 spin columns (GE; 27–5140-01) on a tabletop centrifuge instead of ultra-high speed centrifugation in sucrose cushion. Ribosomal RNA depletion was carried out by using Ribo-Zero Magnetic Kits and this step is moved up to right after the monosomal RNA isolation step and before the Ribosome Protected Fragment gel isolation step.

### Data processing, gene expression quantification, and normalization

For Genotype data, we obtained Genotype VCF files from PsychENCODE BrainGVEX Synapse portal (https://www.synapse.org/#!Synapse:syn5613798). The VCF file includes data from 420 individuals genotyped using a combination of 3 different platforms. Details on genotyping experiments and data processing can be found in our companion manuscripts (*22, 29*). Briefly, data from whole-genome sequencing (WGS), Human PsychChip platform (a custom version of the Illumina Infinium CoreExome-24 v1.1 BeadChip), and Affymetrix GeneChip Mapping 5.0K Array were separately imputed using Minimac3 with Haplotype Reference Consortium (HRC) reference panel and then merged using PLINK with filters for variant call quality, imputation quality, minor allele frequency, and relatedness. Only variants with genotype (or imputed) R^2^ > 0.3, Hardy-Weinberg Equilibrium > 10^-6^ and minor allele frequency (MAF) > 0.01 are kept. The combined and filtered VCF file contains genotype data of 8,108,028 SNPs for 420 individuals. We further evaluated potential effects of additional confounding factors on genotype data (Fig. S21).

For RNA-Seq data, we obtained raw reads in FASTQ format from the PsychENCODE BrainGVEX project (https://www.synapse.org/#!Synapse:syn5613798). We then process the raw reads by removing the adapter sequence using cutadapt (v1.12) code “cutadapt –minimum- length=5 –quality-base=33 -q 30 -a AGATCGGAAGAGCACACGTCTGAACTCCAGTCA -A AGATCGGAAGAGCGTCGTGTAGGGAAAGAGTGT”. The cutadapt processed reads were then mapped onto GENCODE Release 19 (GRCh37.p13) genome with transcript annotations from the corresponding version of GTF files using STAR (v2.4.2a). We used RSEM software (v1.2.29) to quantify read counts for each gene (*56*). The cpm function in the R package “limma” was used to calculate the log-transformed counts per million (CPM). We filtered out genes with CPM < 1 in more than 75% of the samples and samples with network connectivity (*57*) z score < -5 (Fig. S22A), which resulted in a total of 17,207 genes from 426 individuals in the quantification table. We then used normalize.quantiles function in the R package “preprocessCore” (*58*) to normalize expression level for each sample. DRAMS software was used to detect and correct mixed-up samples (*59*), which resulted in a final count of 423 individuals.

For ribo-seq data, we used cutadapt (v1.12) to remove adapter sequence from raw reads with code “cutadapt -a AGATCGGAAGAGCACACGTCT –quality-base=33 –minimum-length=25 – discard-untrimmed”. The processed reads were then mapped against a FASTA file of rRNA, tRNA, and snoRNA sequence downloaded from NCBI using bowtie2 (*60*) to filter out uninformative reads. The filtered reads were mapped to GENCODE Release 19 (GRCh37.p13) genome with corresponding transcript annotation GTF file using STAR (v2.4.2a). The uniquely mapped reads, as defined by the “NH:i:1” flag of the alignment files, were kept for subsequent analysis. We used the featureCounts function in the R package “subreads” (*61*) to calculate gene level read counts for ribosome occupancy. The cpm function in the R package “limma” was used to calculate log-transformed CPM value. We filtered out genes with CPM < 1 in more than 75% of the samples and samples with network connectivity (*57*) z score < -3.5 (Fig. S22B), which resulted in a total of 15,171 genes quantitated from 209 individuals in the quantification table. We then used the normalize.quantiles function in the R package “preprocessCore” (*58*) to normalize ribosome occupancy level for each sample. DRAMS software was used to detect and correct mixed-up samples (*59*), which resulted in a final count of 199 individuals. For quantitative mass spectrometry data, we obtained protein quantification table from the PsychENCODE BrainGVEX Synapse portal (https://www.synapse.org/#!Synapse:syn5613798). This table includes abundance quantification for 11,572 proteins from 268 individuals. The MS/MS raw data were searched against a composite target/decoy human proteome database. The target database contains 83,955 entries, including both canonical proteins and isoforms downloaded from the UniProt database. Since we employed shotgun proteomics, we identified peptides rather than full-length proteins. In cases where a peptide could be mapped to multiple homologous proteins, we followed the rule of parsimony: the peptide was first assigned to the canonical protein. If no canonical form was defined, the peptide was assigned to the protein isoform with the highest uniquely mapped PSM (Peptide-Spectrum Match). For quantification, we summarized all peptides that are mapped to the representative protein. These data processing steps for producing the mass spectrometry quantification table is detailed in Luo et al. (*28*). We further log-transformed protein abundance for each sample. We filtered out genes with log-transformed protein abundance < 1 in more than 75% of the samples and samples with network connectivity (*57*) z score < -6 (Fig. S22C, note that all genes and samples passed the filtering cutoff). We then used the normalize.quantiles function in the R package “preprocessCore” (*58*) to normalize protein level for each sample. DRAMS software was used to detect mixed-up samples (*59*) and found none. We matched protein IDs to gene IDs according to the UCSC database (hg19 version), which resulted in protein level quantifications for 8,330 genes. When matching pQTL signals to genes for comparison with eQTLs/rQTLs, we used the pQTL signals associated with the protein isoform that has the highest median abundance.

### QTL mapping

#### Estimating and adjusting for unwanted factors

We used the R package “PEER” (*62*) to estimate hidden factors for RNA-Seq, ribo-seq, and proteomics data separately. The principle of selecting unwanted hidden factors was to maximize the variance explained with the least number of factors. We identified 30, 29, and 19 hidden factors to remove from RNA-Seq, ribo-seq, and mass spectrometry data, respectively (Fig. S23). For each gene from each omics type, we adjusted the expression level by fitting the selected hidden factors as predictors in a linear model and taking the residuals as the adjusted expression level. The adjusted expression levels were then further centered by mean and scaled by standard deviation.

#### Genotype association tests

We identified *cis*-region expression QTLs (eQTLs), ribosome occupancy QTLs (rQTLs), and protein QTLs (pQTLs) separately using QTLTools (1.3.1) (*63*). Because each gene can encode several protein isoforms, we selected the protein isoform with the highest median abundance as the representative protein. For each gene, we defined *cis*-region as the region ranging from 1Mb upstream of the 5’ end of the gene to 1Mb downstream of the 3’ end of the gene (i.e. the length of the gene body plus 2 Mb in size). We tested all common SNPs (MAF > 0.05) within the *cis*-region for each gene. We performed 10,000 permutations to create null distributions to obtain permutation-based p values and further adjusted permutation-based p values used the beta-approximation approach implemented in QTLTools to calculate empirical p values. For each gene, we selected the most significant SNP to represent the QTL signal (i.e. the variant with the smallest empirical p value) and then calculated a genome-wide FDR using the qvalue function of the R package “qvalue”.

### Annotation and enrichment analyses of QTL variants

We used Ensembl Variant Effect Predictor (VEP) version 105 to annotate QTL SNPs. Each variant was assigned multiple annotations as each variant could map to multiple transcript isoforms of the same gene. We perform three separate analyses with different counting schemes, counting all annotations, counting only most frequent annotations, counting only the annotation with the most severe consequence, to ensure the percentage estimates and conclusions of enrichment are robust.

For genomic feature enrichment analysis, we extracted 5’ UTRs, exons, introns, 3’ UTRs, and intergenic regions from the GENCODE v19 GTF file. The exon regions were extracted directly from entries in the third column of the GTF file (column label: “exon”). Similarly, gene regions were extracted directly from the GTF file. Intron regions were calculated by removing exon regions from gene regions, while intergenic regions were obtained by removing gene regions from the genome. The 5’ UTR and 3’ UTR coordinates were generated using the fiveUTRsByTranscript() and threeUTRsByTranscript() functions from the R package “GenomicFeatures”, respectively. We used bedtools intersect -wa to identify QTL SNPs overlapping each genomic feature. We used Fisher’s exact test to evaluate statistical significance of QTL SNP enrichment in each genomic feature. For genic features (e.g., 5’ UTRs, exons, introns, 3’ UTRs) total genic counts were used as the background; for intergenic features, total cis-region (i.e., mapping window) counts were used as the background.

### Estimating mediated heritability of each omics

We used MESC to estimate the proportion of heritability mediated by different omics separately for four different brain-associated disorders (*33*). In the first step, we calculated the overall expression score using the unwanted-factor-adjusted-expression data (see QTL mapping section) as the individual-level gene expression data and the corresponding BrainGVEX genotype data. We used the 1000 genome phase 3 genotype data to calculate LD scores in order to match BrainGVEX genotype data to GWAS genotype data. In the second step, we used the overall expression score calculated in the previous step and GWAS summary statistics to estimate heritability mediated by each omics.

### Identifying omics-specific QTL

#### SNP-based approach

We used QTLTools to identify omics-specific QTL variants following the same procedure as standard QTL mapping, which is described in the above section, except that the gene expression quantification for the omics-type of interest was adjusted by regressing out signals from the other two omics types. The regression was achieved by including peer adjusted normalized quantification of the other two omics types as covariates in QTLTools. For each gene, we selected the SNP with the most significant genotype-expression association as the candidate omics-specific QTL SNP and used Benjamini-Hochberg procedure to control for false discovery rates. To further remove false positives resulted from ineffective removal of signals from the other two omics types, likely due to poor linear model fit, we removed candidate omics-specific QTLs with nominal significant association (p values < 0.01) with any of the other two omics types.

#### Gene-based PrediXcan approach

Building prediction models: We used the PrediXcan software (*13*) to separately build gene expression prediction models based on RNA-Seq, ribo-seq, and quantitative mass spectrometry data. We built these models for each gene for each omics type with and without regressing out signals from the other two omics types (i.e. for each gene using data of each omics type we build both a standard model and a residual model). For each model, an Elastic Net algorithm was used for feature selection from SNPs located within the *cis*-region defined for each gene (i.e. gene body +/- 1Mb flanking regions) based on results from a ten-fold cross-validation. Weights were produced for every selected SNP of each gene, which were used in the prediction models. For each model, we calculated the Pearson correlation between the predicted and the observed gene expression (Cross-validation R, R_cv_), which was considered as a metric for model prediction accuracy. Unless otherwise specified, we consider models with R_cv_^2^ above the PrediXcan default 0.01 threshold as models describing genes with significant genetic contribution to expression regulation (*13*). We produced prediction models for the unified set of 7,458 genes from 185 samples.

Identifying and categorizing genes containing omics-specific genetic effects: We defined genes with significant residual model as candidate genes containing omics-specific QTL signals using a high R_cv_^2^ cutoff of 0.1 in conjunction with a FDR cutoff of 0.05 for the model fit. Of these candidate genes we further classify them by the model fit results from the other omics types. If no significant standard models were built from any of the two other omics-types, we consider the gene as a predominantly omics-specific QTL gene. For example, for a gene with significant residual model built from RNA-Seq data (eQTL signal), if there are no significant standard model built from either ribo-seq nor proteomics data we consider the gene as a predominantly transcript specific QTL gene (esQTL). For candidate genes that have significant model built from any of the other two omics types we consider these candidate genes as potentially harboring multiple QTL signals from different omics types (either omics-specific or shared QTL effects). To separate out independent signals from each gene we used SuSiE to identify 95% confidence sets representing each individual signals. To categorize these independent signals as shared or omics-specific we then perform *coloc* analysis with data from the other omics-types for each confidence set using SNPs in modest LD (squared correlation > 0.01) with the confidence set of interest to ensure sufficient background SNPs and excluding variants that are in linkage (squared correlation > 0.1) with any other confidence sets. For each confidence set we also performed *coloc* analysis with brain-associated disorder GWAS, in order to identify QTL signals driving disease risks among these confidence sets. See the next section for details on signal colocalization analysis using *coloc*.

### Colocalization

We used *coloc* (*11*) to detect signal colocalization between brain-associated disorder GWAS and each QTL type at the cis-region of each QTL gene. For each QTL gene, for all common SNPs (MAF > 0.05) within the cis-region, we use QTL effects and GWAS summary statistics of brain associated disorders as input for *coloc* analysis. To estimate QTL effects, we calculated, from a linear model fit using SNP genotype as the predictor for gene quantifications, the predictor regression coefficient and its corresponding square of standard error. We then used the coloc.abf function in the R package “*coloc*” (*11*) to calculate the posterior probability of each hypothesis using the default prior. We use posterior probability of 70% for the colocalization hypothesis (i.e. PPH4) as the cutoff for reporting our colocalization findings.

### TWAS identification of SCZ risk genes

We used S-PrediXcan (*34*) to perform gene-level association tests based on the prediction models built using our brain prefrontal cortex multi-omics data, which is described in the “*<u>Gene-based PrediXcan approach</u>*” section, and the SCZ GWAS summary statistics data from PGC3 (*18*). The association tests were performed separately for each omics. For protein data, we performed omnibus test to incorporate p values of all protein isoforms together to calculate a single p value for the corresponding gene. The family wise error rates for SCZ risk genes were calculated using Bonferroni correction of nominal p values.

### Two-sample Mendelian randomization (MR)

To identify causal relationships between each omics type and SCZ we used MR analysis. Here we used fine mapped QTL SNPs as instruments, gene expression quantification at each omics type as exposure, and SCZ GWAS signal as outcome.

More specifically, we took the following steps to test for causal relationship between gene regulation at each omics level and SCZ:

Step 1: Here we used two methods for selecting instrument SNPs: an LD clumping approach and a SuSiE approach. For the LD clumping approach, for each gene of each QTL type, we used PLINK with “–clump-kb 1000 –clump-r2 0.5” parameters and a QTL nominal p value cutoff to define instrument SNPs from separate QTL signals. Since our goal is to test as many risk genes as we reasonably can, we used a dynamic (greedy) approach in defining the QTL p value cutoff for instrument SNP selection. We first set a stringent p value cutoff at 10^-4^, which identified the majority of our instrument SNPs. For genes that don’t have at least 3 instrument SNPs identified we incrementally relaxed the p value cutoff first to 10^-3^ and then to 10^-2^. See Supplemental Table 15 for details on the p value cutoffs used for each omics type by risk gene combination.

Independent of the above mentioned, we performed a separate MR analysis using LD clumping with r^2^ < 0.01 to ensure independence between instrument SNPs and a p value cutoff of 0.05. For the SuSiE approach, we used susie_rss() function with default parameters in the R package “SuSiE” (*47*).

Step 2: We then used the mr_steiger() function in the R package “TwoSampleMR” to test the causal direction of selected instrument SNPs. For genes failing the Steiger test, we applied the steiger_filtering() function to verify the direction of each instrument SNP and excluded those with incorrect directions, ensuring the explained variance by the instrument SNPs was greater for the exposure than for the outcome.

Step 3: For each gene, we used harmonise_data() function in R package “TwoSampleMR” (*64*) to harmonize QTLs of each omics type and SCZ GWAS SNPs to be in the same direction (i.e. effect relative to the same allele).

Step 4: We then performed two-sample MR for each gene by omics type combination separately. Two-sample MR analysis was done using the R package “MendelianRandomization” (*65*). We used mr_ivw(data, correl = TRUE) function to conduct generalized IVW and mr_egger(data,correl = TRUE) function to conduct generalized Egger (see next step). Generalized IVW accounts for linkage between instrument SNPs, which enables the MR analysis to use linked instrument SNPs.

Step 5: We used the intercept test (i.e. the Egger method) to test for horizontal pleiotropy, and used predictor coefficient and its corresponding p value from IVW to determine the effect size and significance of causal effects for each omics type on SCZ.

Step 6: We used Benjamini-Hochberg control procedure to adjust the nominal p values from IVW for multiple testing.

Step 7: We define a gene by omics type combination as causal for SCZ if the causal effect test passed the significance cutoff of FDR < 0.1 and the Egger intercept test passed the intercept p value > 0.05 cutoff.

### One-sample Mendelian randomization

For one-sample MR we used Two-stage predictor substitution (TSPS) approach, which is a method to find causal relationships between omics types. We performed the following analysis in two iterations, both following the direction of genetic information flow. In the first iteration, we tested causal relationships between transcript level and ribosome occupancy level (i.e. mRNA -> ribosome occupancy). In the second iteration, we tested causal relationships between ribosome occupancy level and protein level (i.e. ribosome occupancy -> protein).

Step 1: We used LD clumping for instrument SNP selection as described in the previous section. Here we tested two pathways (mRNA -> ribosome occupancy and ribosome occupancy -> protein). For mRNA -> ribosome occupancy, we used the SNPs of eQTL with p value < 10^-4^ and R^2^ < 0.5 from LD. In order to test as many risk genes as we reasonably can, the same dynamic p value cutoff approach (i.e. the same as the above described for two-sample MR) was used for instrument SNP selection. See Supplemental Table 17 for details on the p value cutoffs used for each risk gene.

Step 2: We then used the mr_steiger() function in the R package “TwoSampleMR” to test the causal direction of selected instrument SNPs. For genes failing the Steiger test, we applied the steiger_filtering() function to verify the direction of each instrument SNP and excluded those with incorrect directions, ensuring the explained variance by the instrument SNPs was greater for the exposure than for the outcome.

Step 3: We then combined genotype and quantification data of the relevant omics types into a dataframe: dat_mr_ for mRNA -> ribosome occupancy pathway, dat_rp_ for ribosome occupancy -> protein pathway.

Step 4: For each gene, we then set formula in R package “OneSampleMR” (*66*): tsps (“ribosome occupancy ∼ mRNA | SNP_1_+SNP_2_+SNP_3_+…+SNP_n_”, data = dat_mr_) and tsps (“protein ∼ ribosome occupancy | SNP_1_+SNP_2_+SNP_3_+…+SNP_m_”, data = dat_rp_).

Step 5: We used the summary() function to get the slope (i.e. the regression coefficient) and the corresponding p value of the slope from each tsps object.

Step 6: We used Benjamini-Hochberg control procedure to adjust the p value of the slope for multiple testing.

Step 7: We defined a relationship for each gene by pathway combination as causal if the test results passed the significance cutoff of FDR < 0.1.

Step 8: We also used R package “ivreg” with the same dataset and formula as the above described for tsps objects to calculate F-statistics for each gene. The calculated F-statistics was used to check instrument strength.

## Data availability

All raw data from psychENCODE BrainGVEX project are available through https://www.synapse.org/#!Synapse:syn5613798.

## Code availability

All code, and processed data used in the analyses are available at https://github.com/qiumanL/multi-omics-qtl.

## Acknowledgments

We thank the three anonymous reviewers for their valuable comments, which resulted in substantial improvement of the manuscript. We thank Majd Alsayed, Miguel Brown, Dominic Fitzgerald, Amber Thomas, and Mimi Brown for their assistance with RNA-Seq data production and processing. We thank Richard Kopp at SUNY Upstate Medical University for his help with wordsmithing. We thank all of the donors and their relatives participated in the brain collections at Stanley Medical Research Institute and Banner Sun Health Research Institute. We thank Drs. Maree J. Webster and Thomas Beach for their supports to make the tissue available. We thank the High Performance Computing Center of Central South University for partial support of calculation. This project has received funding from National Key Research and Development Program of China (2024YFA1108000), National Natural Science Foundation of China (Grants No. 82022024), SUNY Empire Innovation Program, National Institutes of Health grants (U01MH122591, U01MH116489, R01MH110920, R01MH126459, U01MH103340, R01MH109715, R01MH110555, R21MH129817, R01GM139980).

## Author contributions

CL, KPW, JP, SHW, BL, XW, CC were responsible for Conceptualization. AWS, LC, DZ, FW, MX, MN, DP, YJW, RV, CZ, KG, GG were responsible for sample collection, DNA extraction, mRNA extraction, mRNA binding with ribosome extraction, protein extraction, sequencing, mass spectrum and raw data processing. SHW, QL, YJ, CC, CY, AWS were responsible for investigation of analysis. QL was responsible for visualization of results. SHW, CL, CC, BL, KPW were responsible for supervision. SHW, QL, YJ, CL were responsible for original draft writing. SHW, CL, CC, QL were responsible for review and editing.

