## Supplemental materials for "The impact of common variants on gene expression in the human brain: from RNA to protein to schizophrenia risk"

### Supplementary Materials

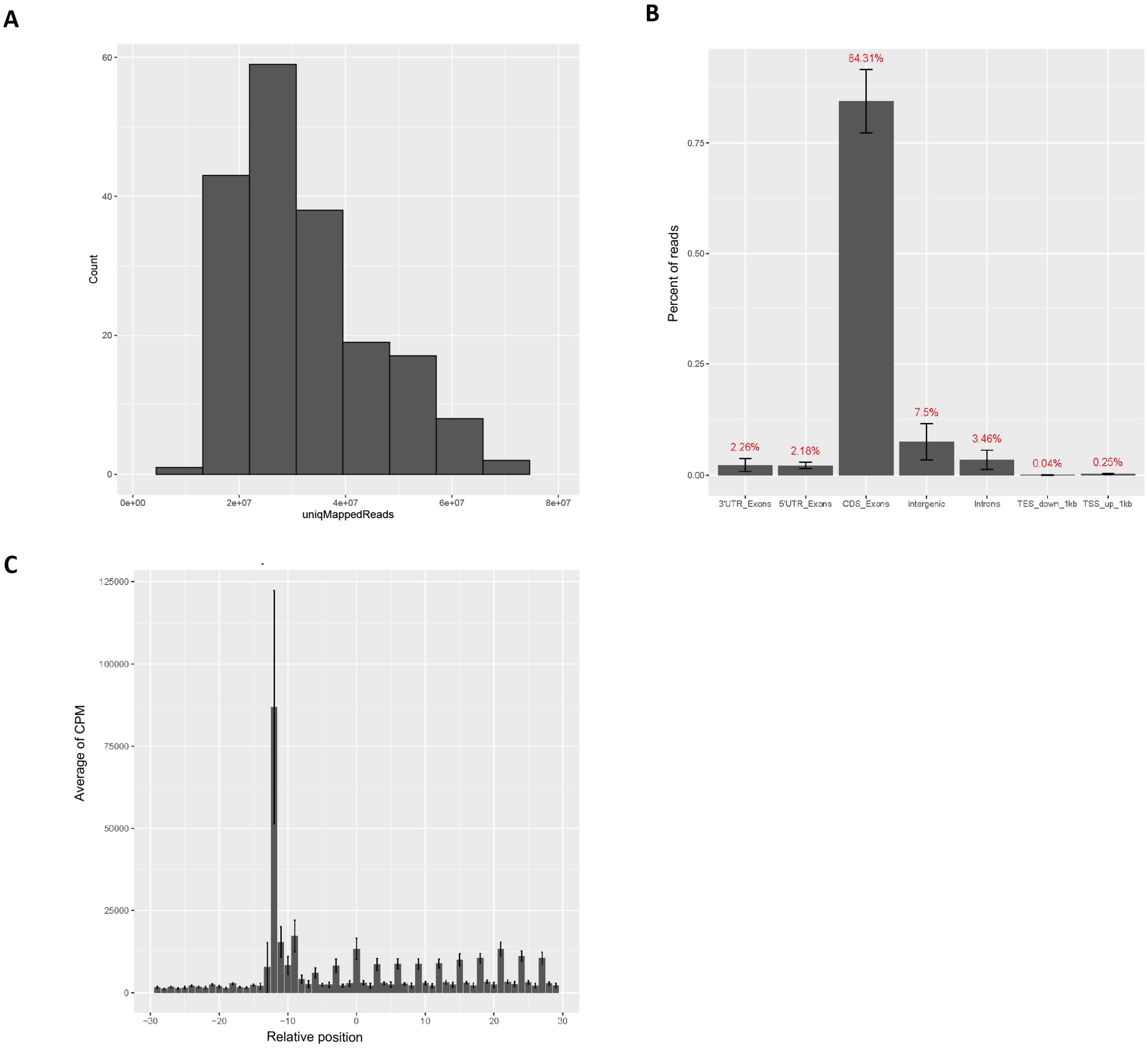


**Fig. S1. Summary statistics presenting key metrics for evaluating ribo-seq data quality. (A)** A histogram summarizing the distribution of the number of uniquely mapped ribo-seq reads collected across samples. **(B)** A bar plot summarizing the proportion of ribo-seq reads mapped to each genic features (TSS: Transcription Start Site, TES: Transcription End Site). **(C)** Aggregated Subcodon periodicity pattern at the +/- 30 nt window around GENCODE annotated translation initiation sites. Ribo-seq read counts were aggregated across genes according to the distance between the 5’ end of each mapped read and the annotated translation initiation sites. X-axis:

relative distance to the translation initiation site, one nucleotide per bar. Y-axis: number of reads aggregated at each relative position. Error bars represent standard deviation across samples. Note the 13 nt offset of the start codon enrichment peak upstream to the annotated translation initiation sites, this offset reflects the physical distance between the 5' edge of the ribosome to the A site of the actively engaged ribosome.

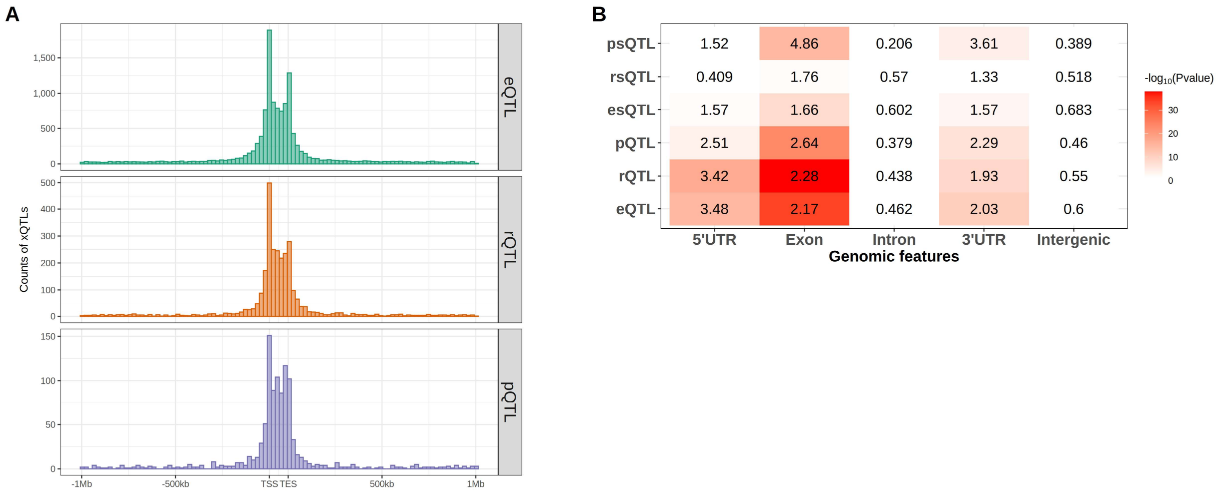


**Fig. S2. Enrichment of genomic features and variant effects for eQTL, rQTL, pQTL, esQTL, rsQTL and psQTL. (A)** Histograms summarizing the distance of each QTL to its corresponding gene and genic features. The distance distribution within the gene body is scaled to the gene length. TSS: Transcription Start Site. TES: Transcription End Site. **(B)** An enrichment heatmap of genomic features. Numbers indicate the odds ratio of enrichment for each genomic feature based on Fisher's test compared to the background. Background definitions are as follows: for exon, intron, 5' UTR, and 3' UTR, the background is genic features (i.e. the sum of exon, intron, 5' UTR and 3' UTR). For intergenic regions, the background is all genomic features (i.e. all variants within the QTL mapping window). Colors represent -$\log_{10}$(P value) of enrichment for each genomic feature from Fisher's exact tests.

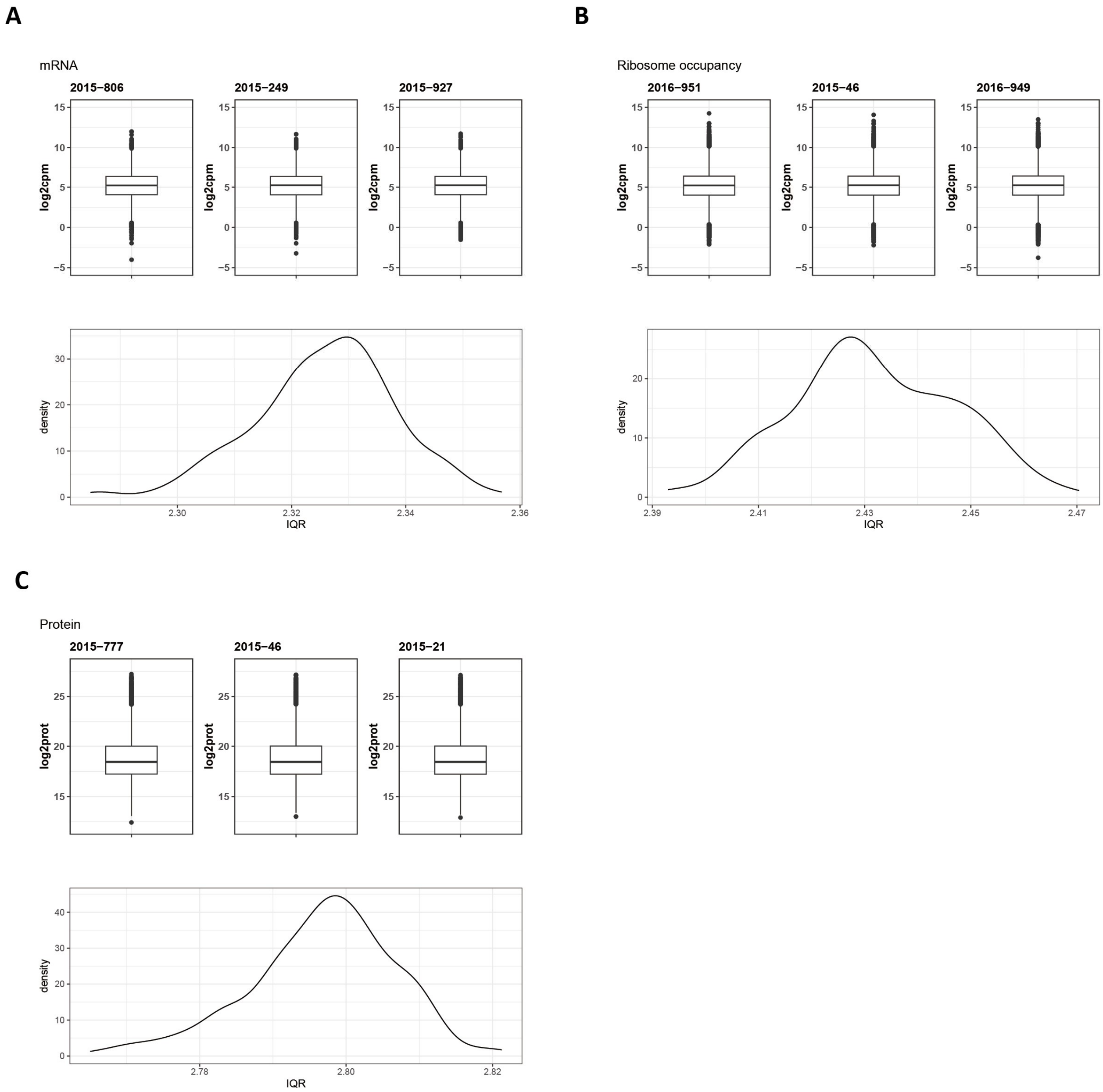


**Fig. S3. Dynamic range of gene quantification for each omics.** (**A**) RNA-Seq. (**B**) Ribo-seq. (**C**) Quantitative mass spectrometry. The density plots present the distribution of interquartile range (IQR) of quantification across all genes across samples. The boxplots showing range of expression level, from left to right, correspond to individual samples in the first quartile, median, and third quartile of the IQR.

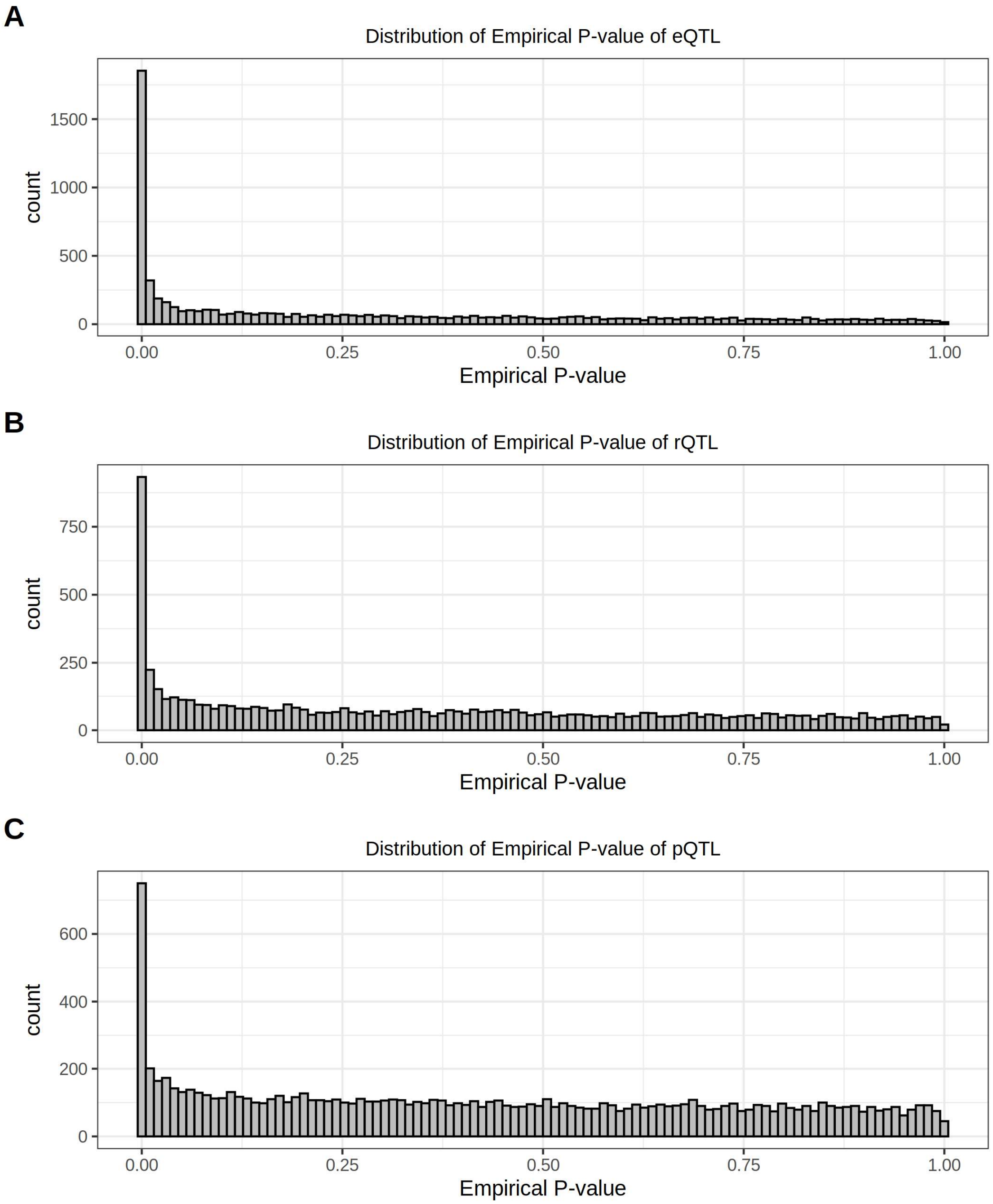


**Fig. S4. Histograms of QTL mapping permutation p values. (A)** eQTL. **(B)** rQTL. **(C)** pQTL.

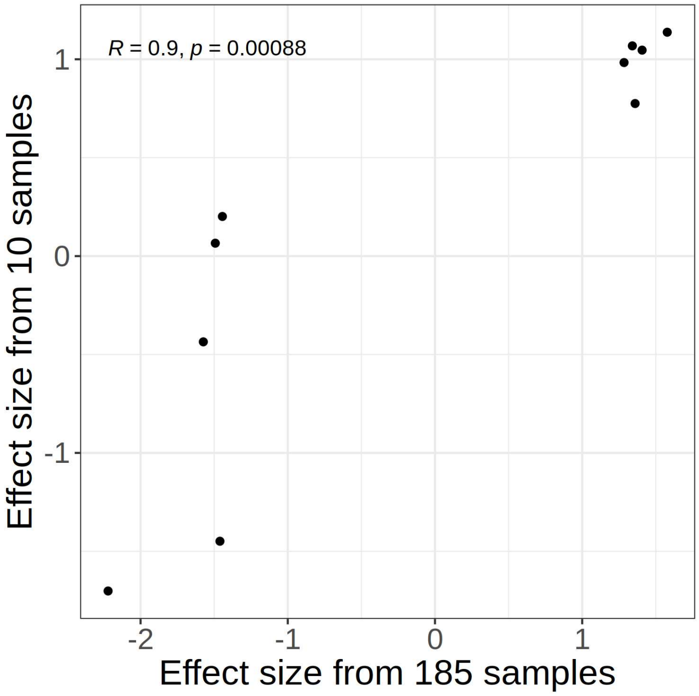


**Fig. S5. Replication of rQTLs discoveries.** A scatter plot comparing the rQTL effect size from the replication tests to the original discovery. We replicated our findings using a small cohort of 10 additional individuals. Given the small sample size in the replication cohort we have rather limited power to detect and in many cases don't have enough minor alleles to perform the analysis. We therefore focus our analysis on large effect size rQTLs that have at least 3 copies of the minor alleles in the replication cohort. Of the top 10 rQTL discoveries (ranked by effect size), 8 show the same direction of effects in the replication cohort, 6 of these 8 rQTL replicated at 10% FDR (one-sided test). The replication rate is clearly driven down by sampling error given that the effect size estimates between the discovery and replication tests are strongly correlated (R = 0.9).

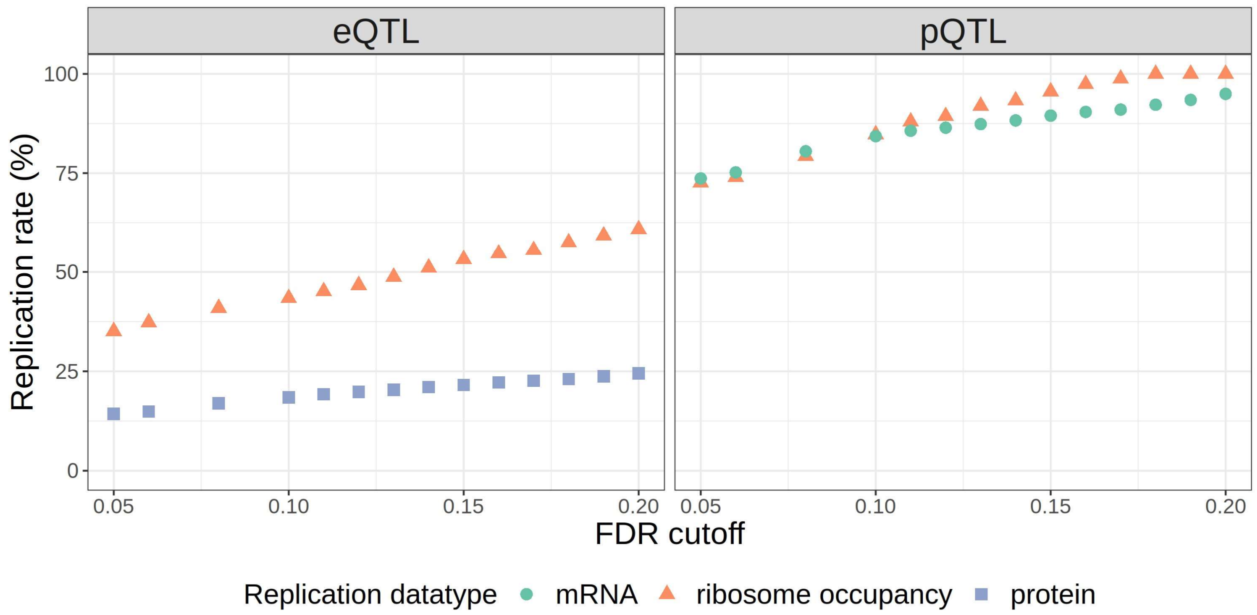


**Fig. S6. Replication rates between QTL types.** The discovery QTL types are labeled at the top. The replication datatypes are color and shape coded with the color and shape key positioned at the bottom of the plots. X-axis: the FDR cutoff used to define QTL replication (i.e. from 5% to 20%). Y-axis: percentage of QTLs replicated.

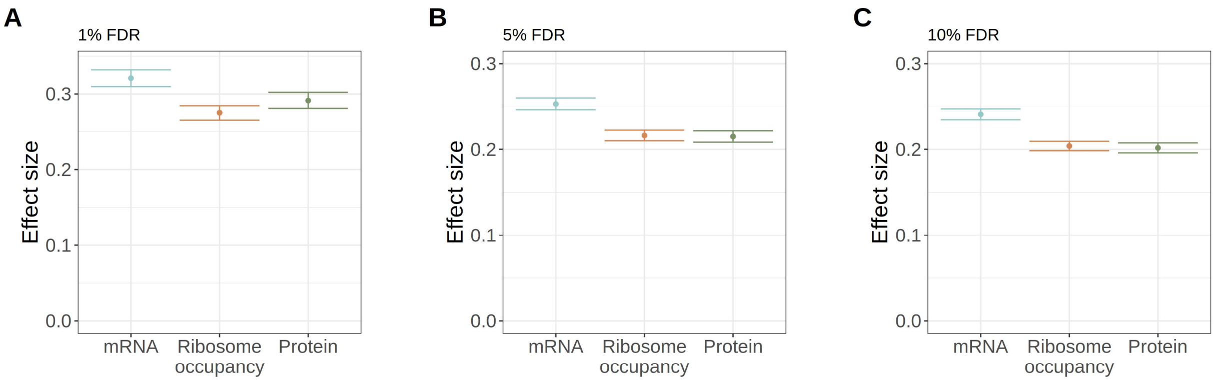


**Fig. S7. Effect size of ROSMAP pQTL variants in BrainGVEX dataset.** Mean +/- standard error of per allele effect on expression in each of the three BrainGVEX omics (i.e. RNA-Seq, ribo-seq, and quantitative mass spectrometry) for pQTLs identified from the ROSMAP dataset at **1%** FDR **(A)**, **5%** FDR **(B)**, and **10%** FDR **(C)**.

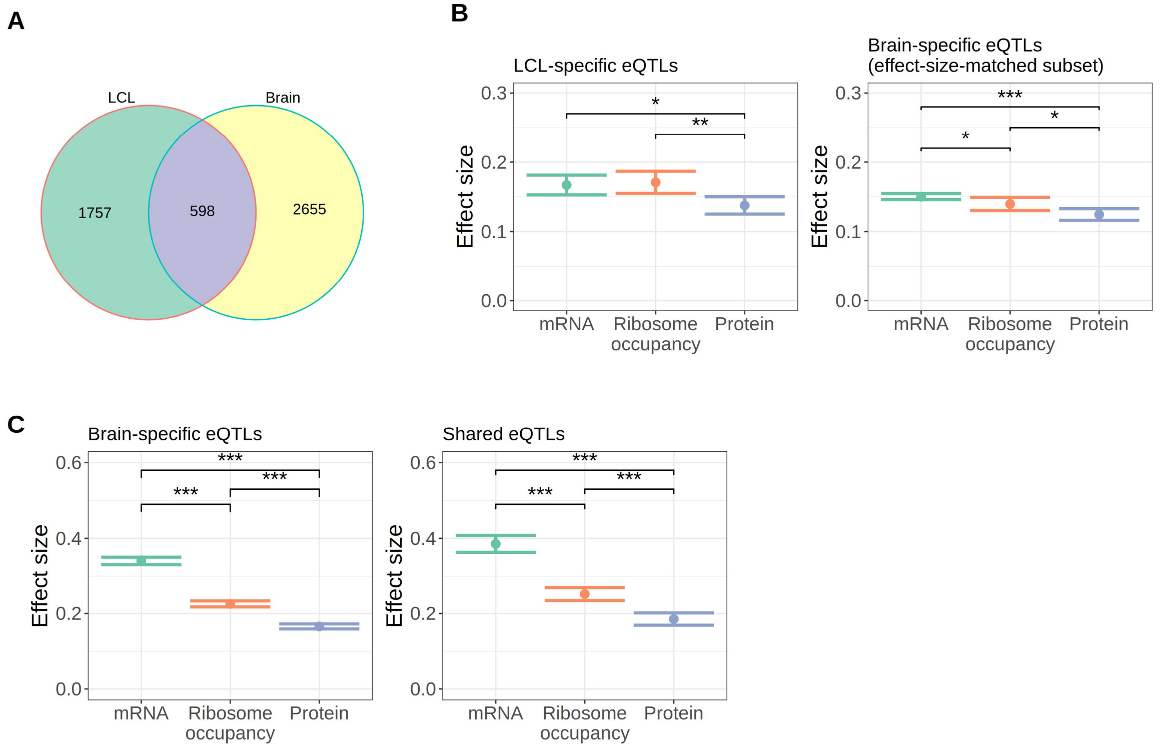


**Fig. S8. Comparisons of QTL mapping results between the Battle et al. LCL study and our BrainGVEX prefrontal cortex study. (A)** A Venn diagram comparing the eGene discoveries between the two studies. **(B)** Comparisons of eQTL effect in BrainGVEX data between LCL-specific eQTLs and a selected subset of brain-specific eQTLs. This subset is selected to match the effect size of eQTLs identified only in the Battle et al. LCL study to enable a fair comparison. **(C)** Comparisons of eQTL effect in BrainGVEX data between brain-specific eQTLs and shared eQTLs. Significance levels for one-sided t-tests for eQTL effect attenuation: $p < 0.05$ *, $p < 0.001$ **, $p < 0.0001$ ***.

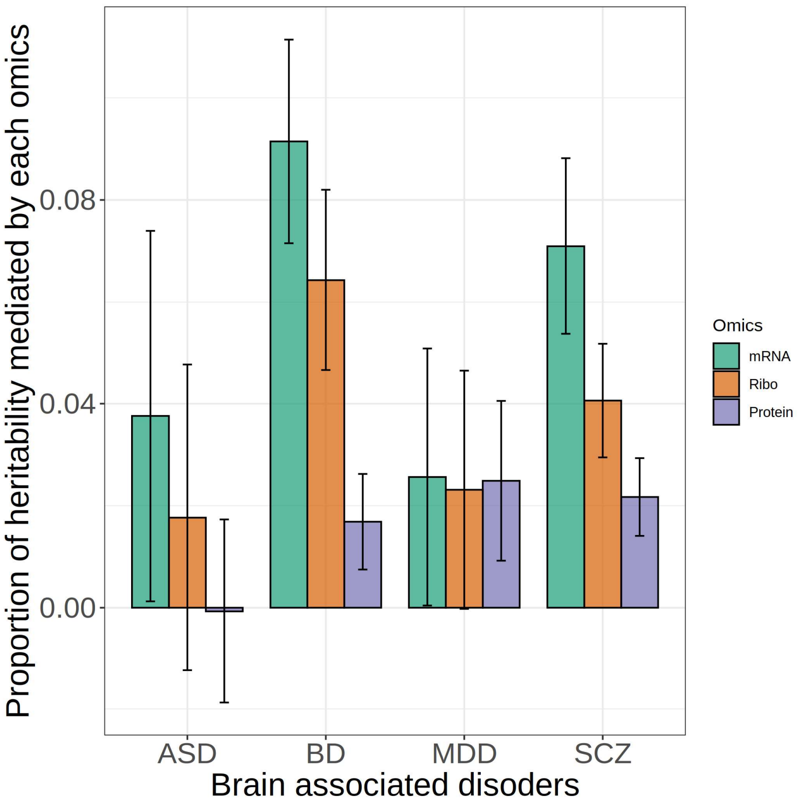


**Fig. S9. Proportion of heritability mediated by gene expression for brain associated complex disorders.** A bar plot presenting the mean and standard error of proportion heritability mediated by each omics for Autism Spectrum Disorder (ASD), Bipolar Disorder (BD), Major Depressive Disorder (MDD) and Schizophrenia (SCZ). Ribo: ribosome occupancy.

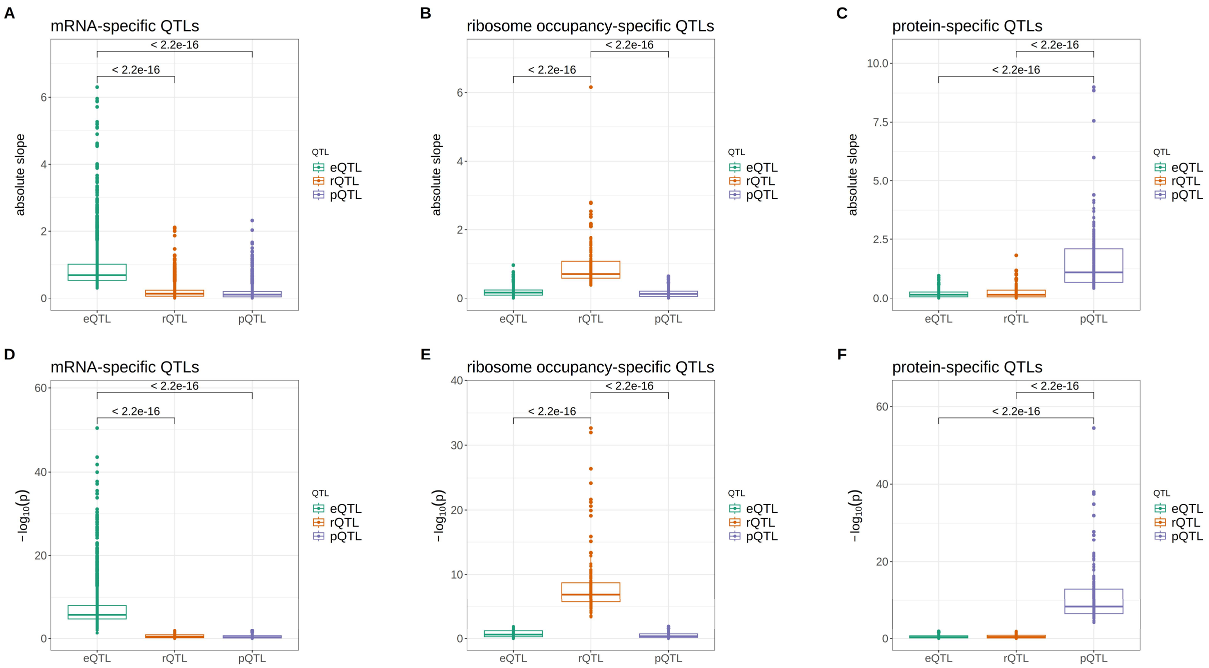


**Fig. S10. Omics-specific QTLs show the expected patterns in effect size and p value across QTL types. (A, B, C)** Boxplots summarizing the QTL effect size and **(D, E, F)** Boxplots summarizing the corresponding -$\log_{10}$(p value).

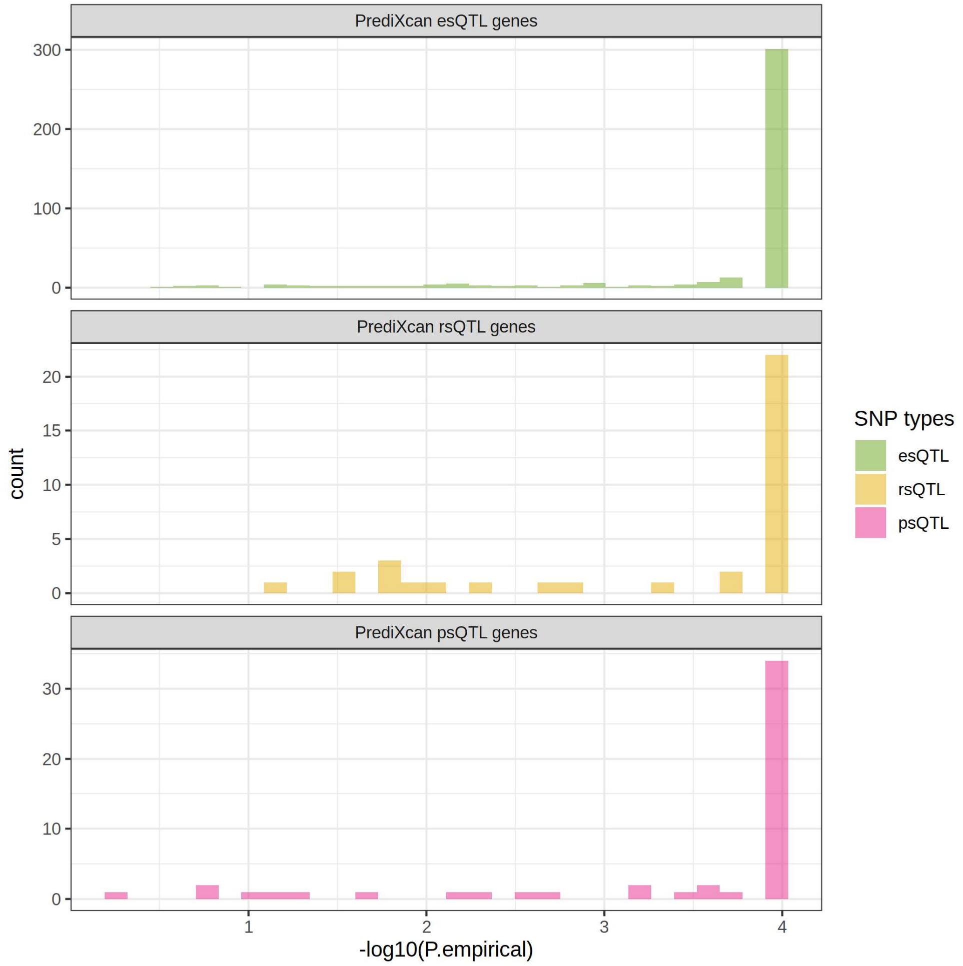


**Fig. S11. Omics-specific QTL genes identified by the gene-based PrediXcan approach are enriched of significant omics-specific QTL variants identified using the SNP based approach.** Histograms summarizing the distribution of -$\log_{10}$(p value) of omics-specific QTL variants identified among omics-specific QTL genes.

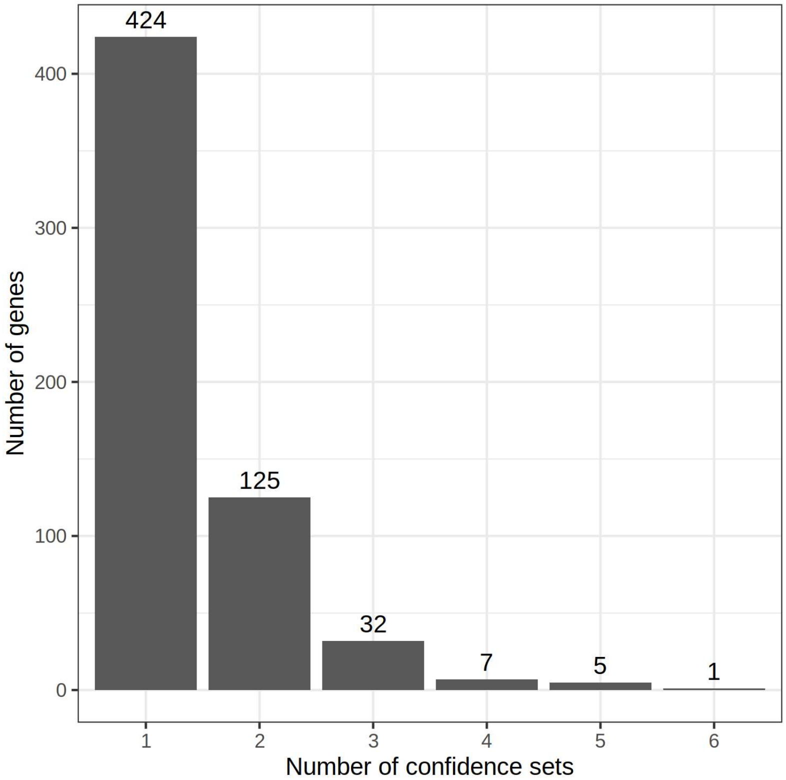


**Fig. S12. Number of SuSiE confidence sets identified for candidate genes that potentially contain both omics-specific and shared QTL signals.** A histogram summarizing the distribution of the number of confidence sets identified per gene.

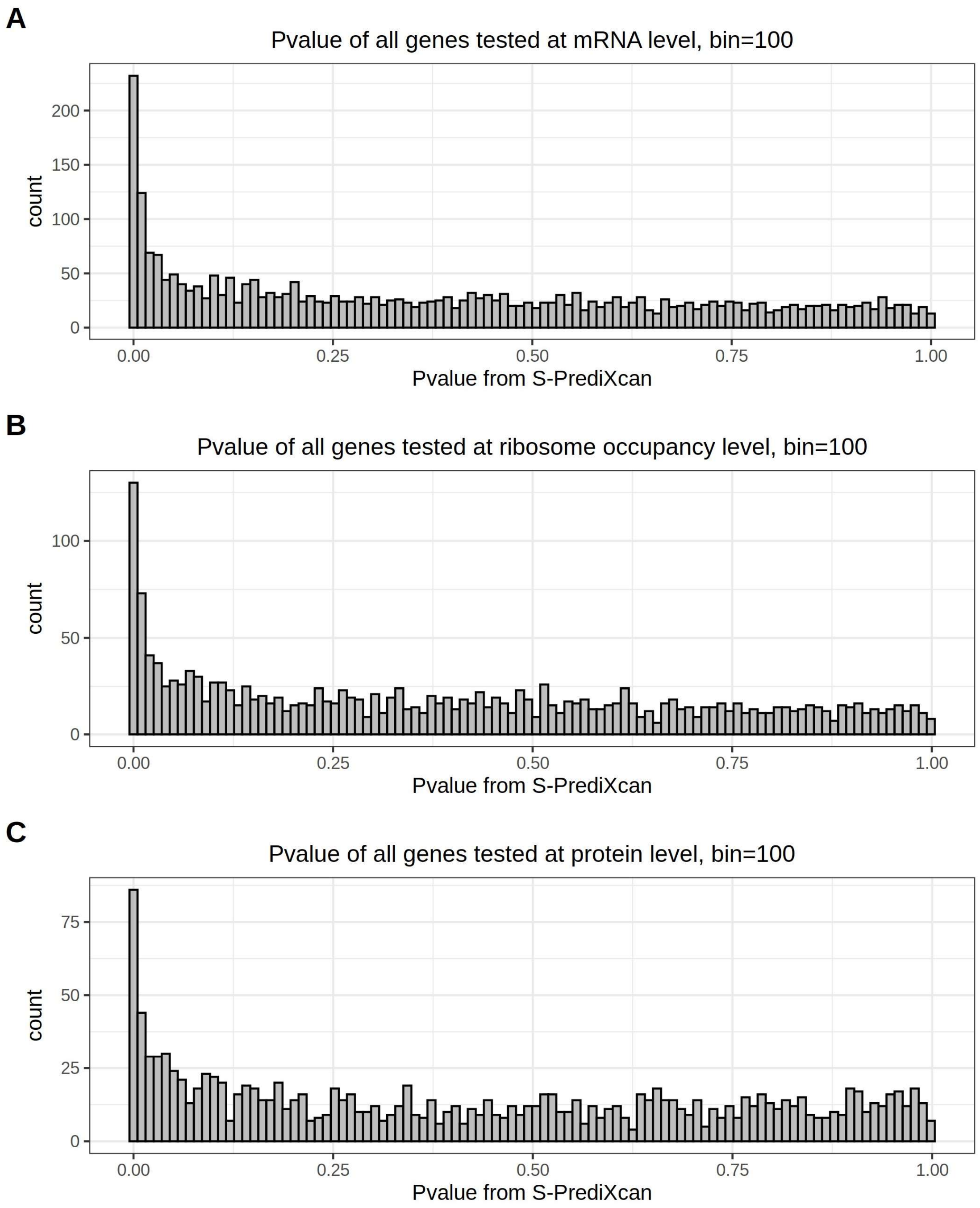


**Fig. S13. Distribution of SCZ risk gene detection p-values from S-PrediXcan analyses.** Results presented in each panel are histograms summarizing p-values from analyses using **(A)** transcript level (RNA-seq), **(B)** ribosome occupancy level (ribo-seq), and **(C)** protein level (quantitative mass spectrometry).

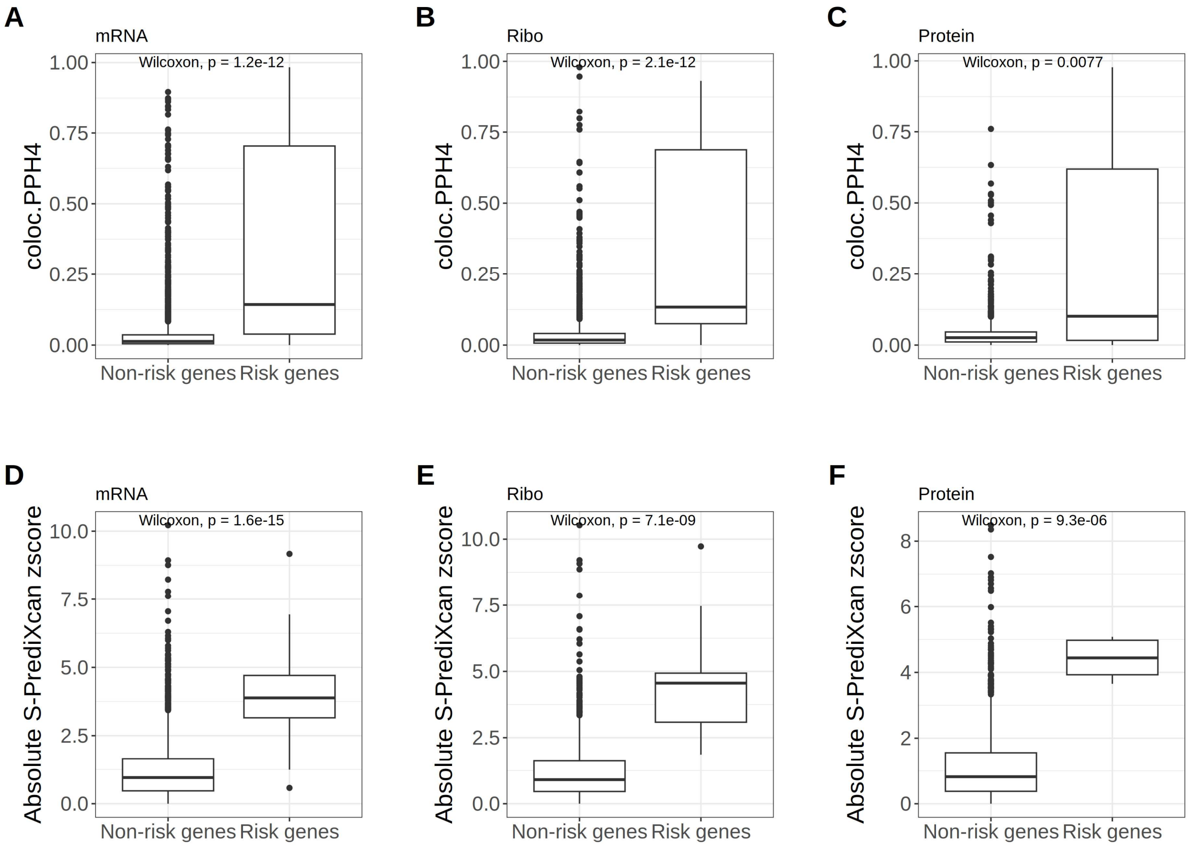


**Fig. S14. Comparison of SCZ risk gene identification results between S-PrediXcan and *coloc*.** Boxplots summarizing *coloc* posterior probability (i.e., PPH4) for SCZ risk genes identified by S-PrediXcan for mRNA (**A**), ribosome occupancy (**B**), and protein (**C**). Boxplots summarizing S-PrediXcan z-scores for SCZ risk genes identified by *coloc* for mRNA (**D**), ribosome occupancy (**E**), and protein (**F**).

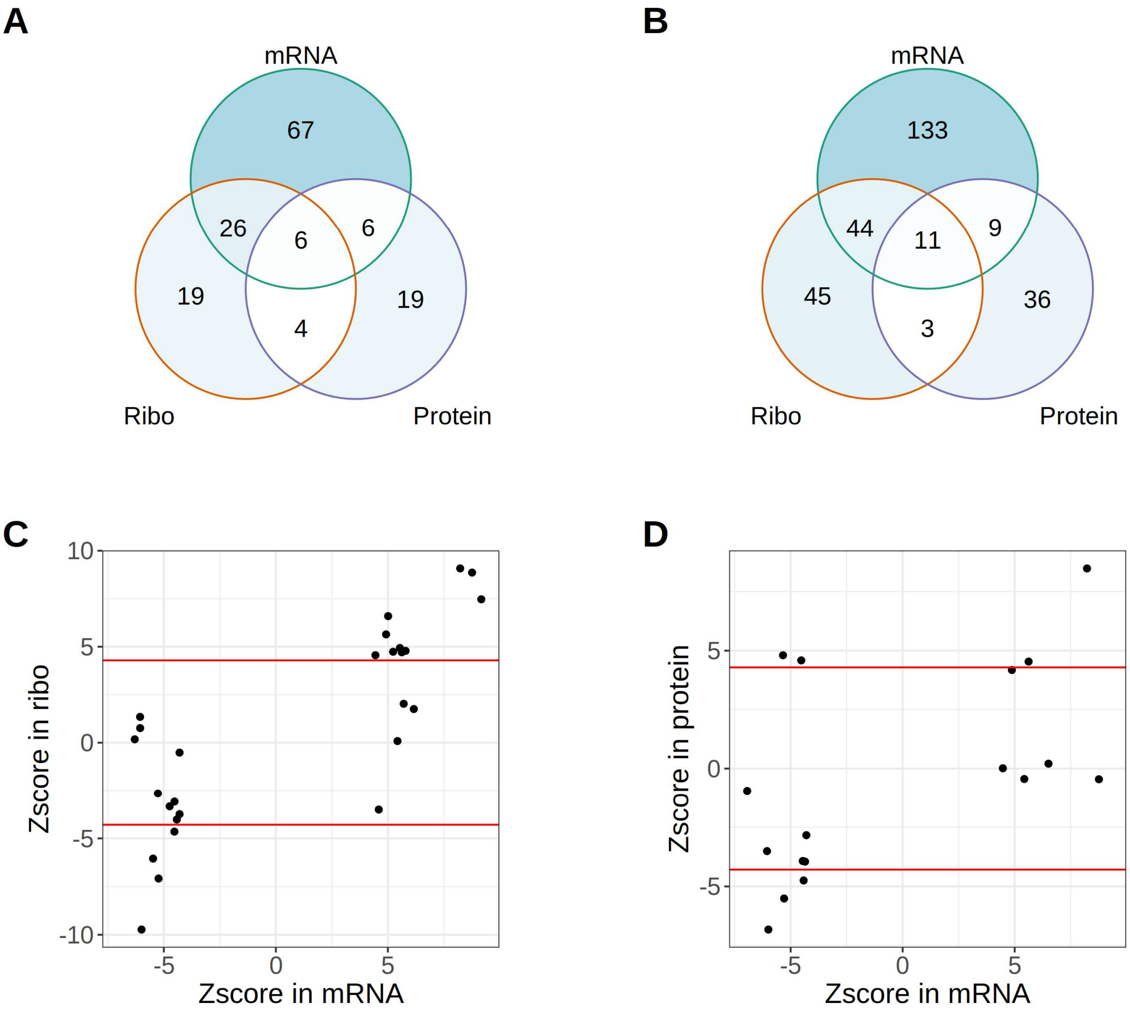


**Fig. S15. A lack of overlap in SCZ risk genes identified between different omics is consistently observed across multiple significance levels. (A)** A Venn diagram of SCZ risk genes identified from each omics at 1% false discovery rate. **(B)** A Venn diagram of SCZ risk genes identified from each omics at 5% false discovery rate. **(C, D)** Scatter plots of S-PrediXcan risk gene z-scores for mRNA risk genes to demonstrate that the limited overlap of risk genes identified between different omics is not simply a result of a specific significance cutoff used. The red lines indicate the z-score corresponding to a family-wise error rate of 5%. Note that not all mRNA risk genes are presented in the scatter plots because for some mRNA risk genes the corresponding z-scores from the other omics are not available likely because of a lack of effect there.

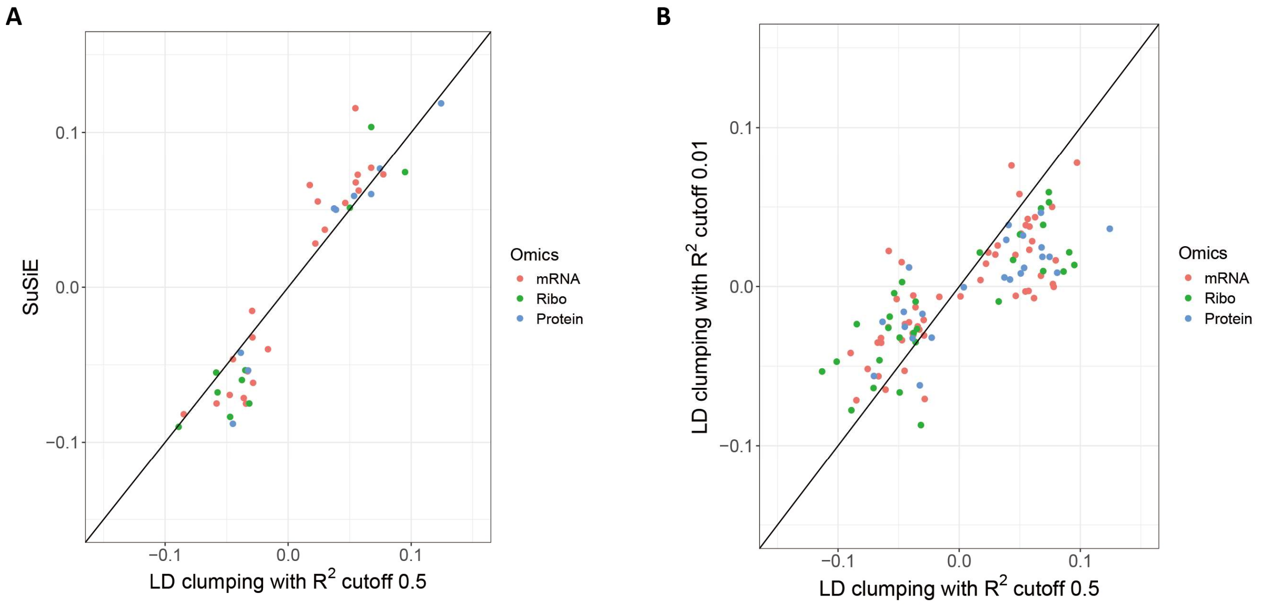


**Fig. S16. Three different methods of selecting instrumental variables were implemented in the two-sample MR analyses for testing causal effects between QTL and SCZ risk genes.** Each datapoint in the scatterplot represents a risk gene, causal effects between QTL and schizophrenia GWAS signals calculated using LD clumped SNPs with $R^2 < 0.5$ as instrumental variable (X-axis) is plotted against parallel analyses using SuSiE fine-mapped SNPs as instrumental variable (Y-axis) **(A)** or using LD clumped SNPs with $R^2 < 0.01$ as instrumental variable **(B)**. Because SuSiE failed to provide instrument SNPs for some SCZ risk loci, there were fewer datapoints shown in (A) comparing to (B). Ribo: ribosome occupancy.

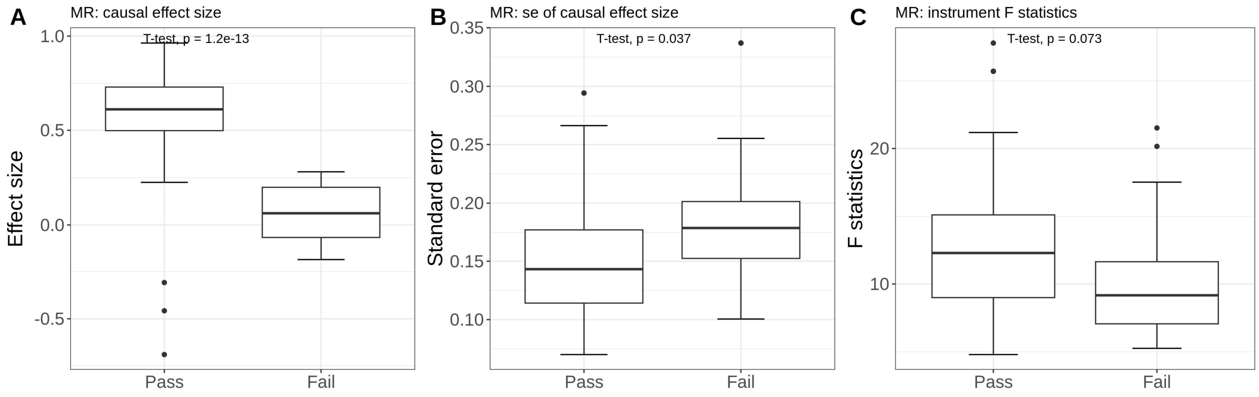


**Fig. S17. Boxplots summarizing key statistics from one-sample-MR tests for the both-passed genes (labeled as Pass) versus the none-passed genes (labeled as Fail). (A)** Coefficients associated with the predictor of the MR tests (i.e., the effect size). **(B)** Standard error of the predictor coefficient. **(C)** F-statistics from the exposure-instrument variable regression.

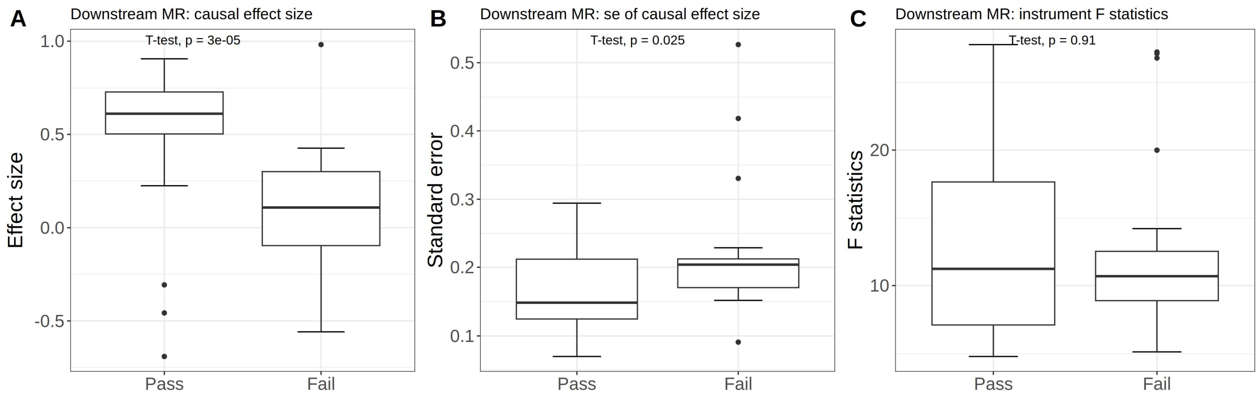


**Fig. S18. Boxplots summarizing key statistics of the downstream pathway from one-sample-MR tests for the both-passed genes (labeled as Pass) versus the upstream-passed genes (labeled as Fail). (A)** Coefficients associated with the predictor of the MR tests (i.e., the effect size). **(B)** Standard error of the predictor coefficient. **(C)** F-statistics from the exposure-instrument variable regression.

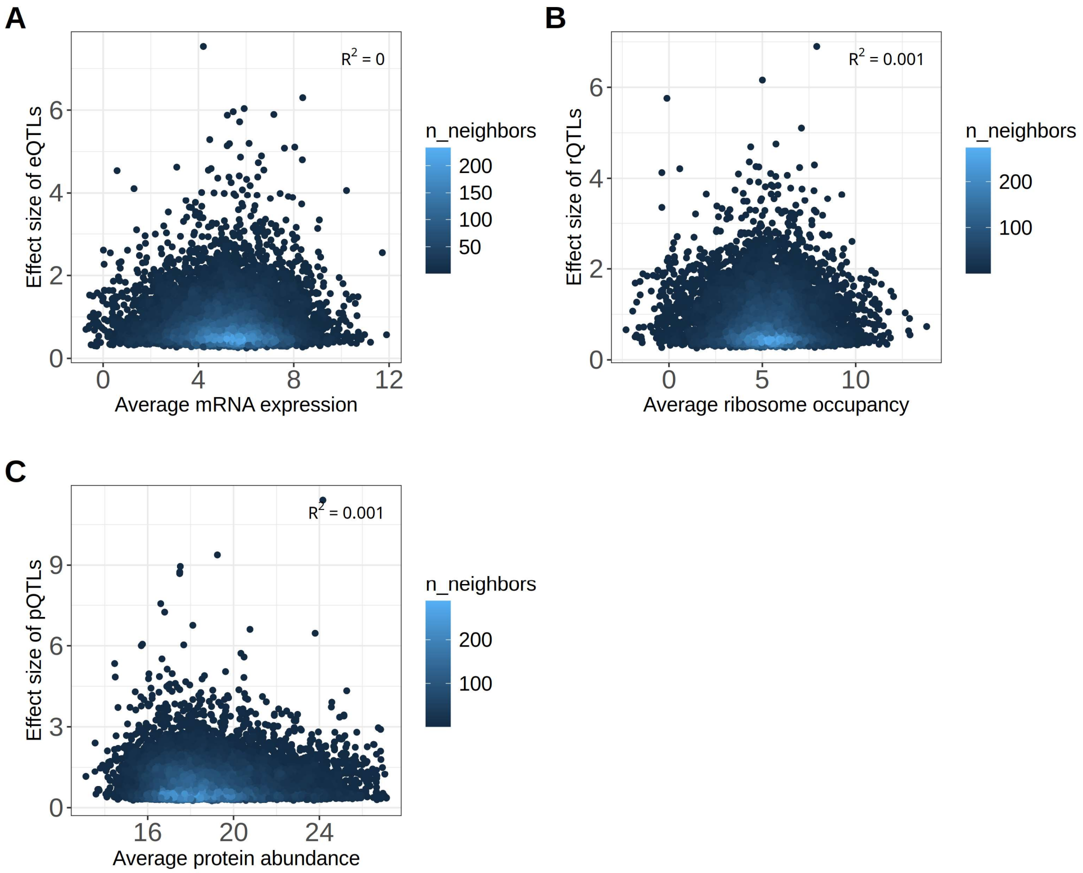


**Fig. S19. Scatter plots of QTL effect size vs. gene quantification for each omics. (A)** eQTL effect size vs. mRNA expression. **(B)** rQTL effect size vs. ribosome occupancy. **(C)** pQTL effect size vs. protein abundance. Color code reflects the density of data points.

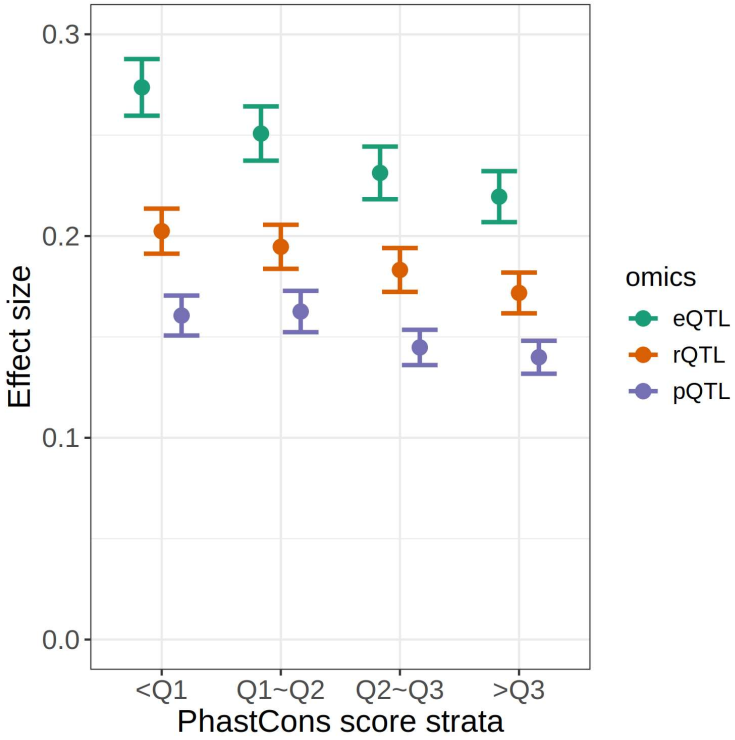


**Fig. S20. QTL effect size across PhastCons score strata.** The higher a gene's PhastCons score, the more conserved it is. Q1, Q2, and Q3 represent the first, the second, and the third quartiles of the PhastCons score, respectively.

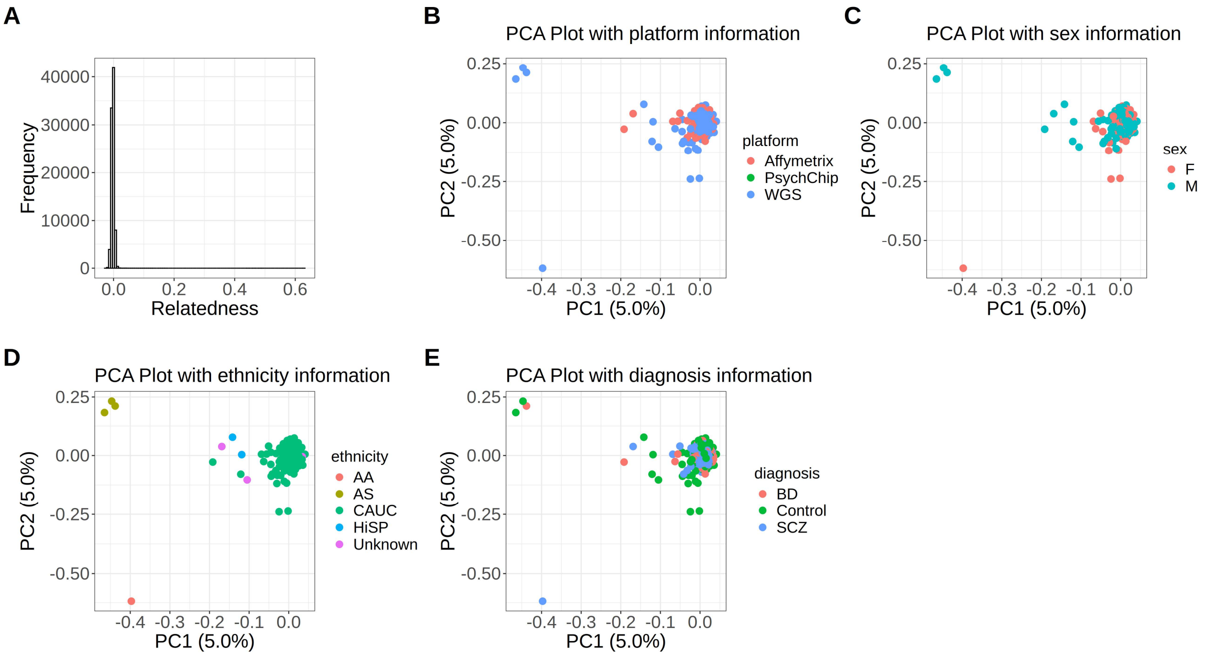


**Fig. S21. Genotype information for 420 individuals from the BrainGVEX cohort. (A)** A histogram summarizing the inter-individual relatedness. **(B, C, D)** Scatter plots visualizing results from Principal Components Analyses (PCA): inter-individual variation in genotype data is projected on to the top two principal components. **(B)** Genotype PCA plot color coded by genotyping platform. Affymetrix and PsychChip are two different genotyping arrays. WGS: Whole Genome Sequencing. **(C)** Genotype PCA plot color coded by sex. F: female. M: male. **(D)** Genotype PCA plot color coded by ethnicity. AA: African American. AS: Asian. CAUC: Caucasian. HISP: Hispanic. **(E)** Genotype PCA plot color coded by psychiatric disorder diagnosis status. BD: bipolar disorder, SCZ: schizophrenia, Control: not diagnosed with psychiatric disorders.

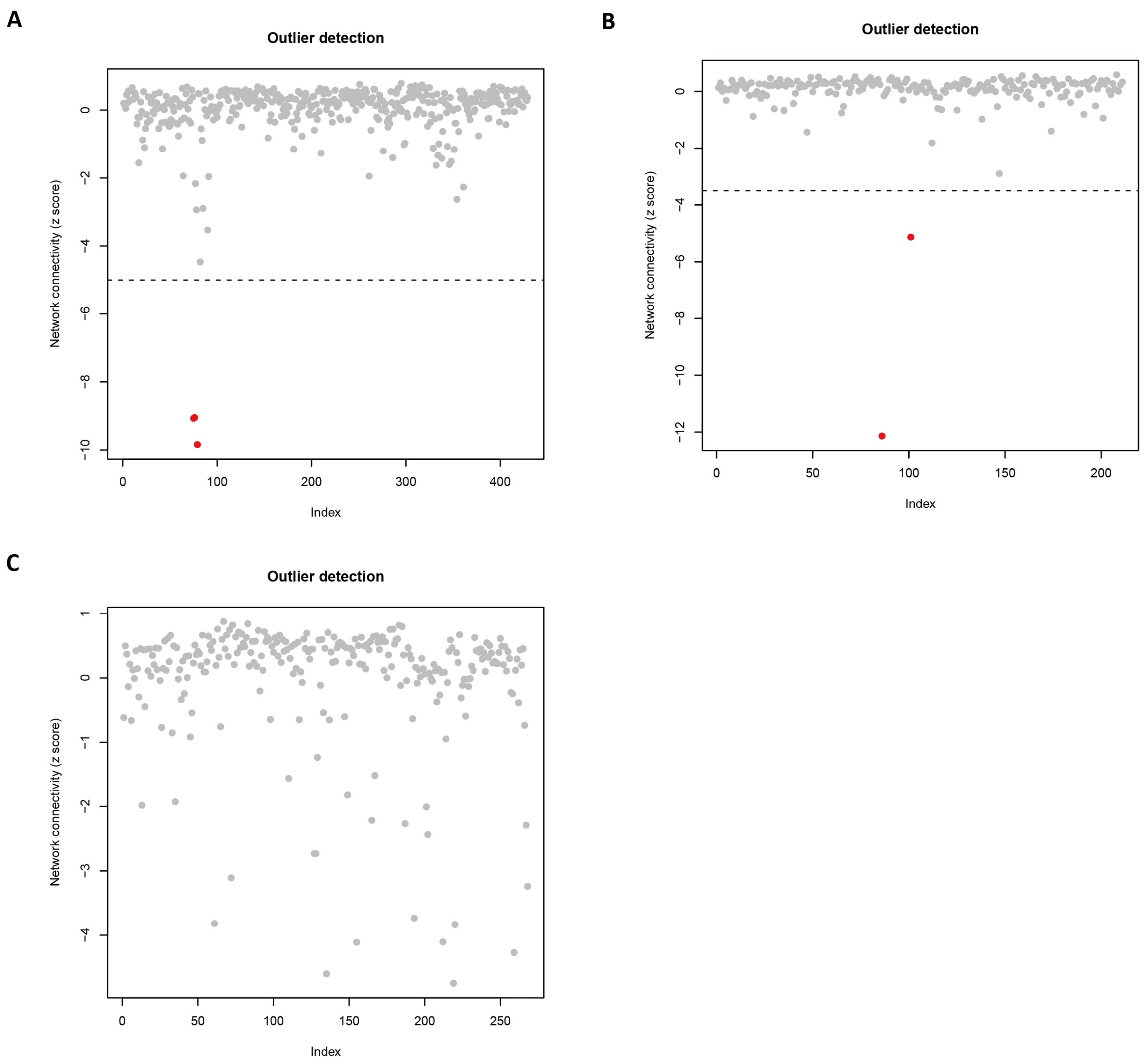


**Fig. S22. Standardized network connectivity calculated across all genes for each sample was used to identify outlier samples.** Network connectivity z score (Y-axis) is shown for each sample (X-axis, ordered by sample ID) in a scatter plot for **(A)** RNA-seq. **(B)** Ribo-seq. (**C**) Quantitative mass spectrometry. The red dots denote the samples excluded from the QTL analysis. The dashed lines denote the z score cutoff.

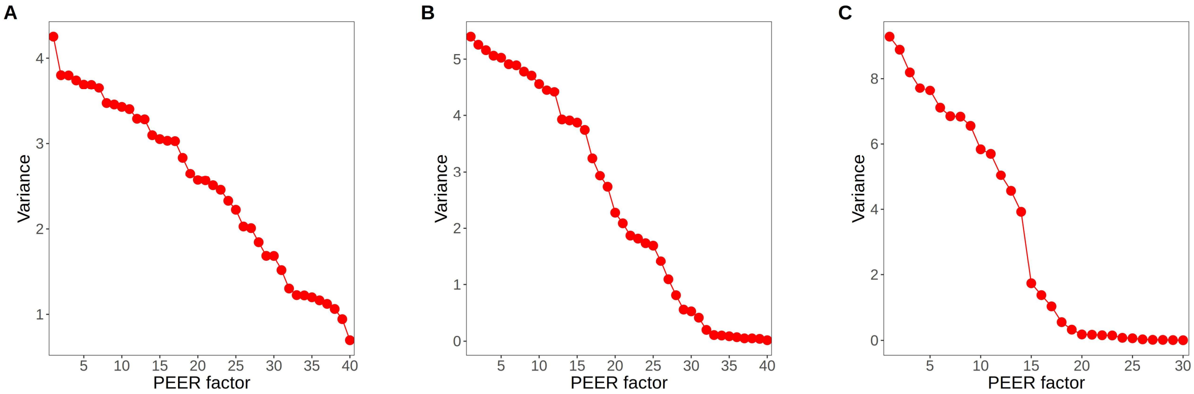


**Fig. S23. Variance in BrainGVEX multi-omics data explained by PEER factors. (A)** RNA-Seq. **(B)** Ribo-seq. **(C)** Quantitative mass spectrometry. PEER: Probabilistic Estimation of Expression Residuals.

#### PsychENCODE Consortium Authors and Affiliations

‡ Schahram Akbarian[1], Alexej Abyzov[2], Nadav Ahituv[3], Dhivya Arasappan[4], Jose Juan Almagro Armenteros[5], Brian Beliveau[6], Jaroslav Bendl[1], Sabina Berretta[7], Rahul Bharadwaj[8], Arjun Bhattacharya[9], Lucy Bicks[9], Kristen Brennand[10], Davide Capauto[10], Frances A. Champagne[4], Tanima Chatterjee[10], Christos Chatzinakos[7], Yuhang Chen[10], Han-Chia Chen[11], Yuyan Cheng[9], Lijun Cheng[12], Andrew Chess[1], Jo-fan Chien[13], Zhiyuan Chu[10], Declan Clarke[10], Ashley Clement[3], Leonardo Collado-Torres[8], Gregory Cooper[14], Gregory Crawford[15], Rujia Dai[16], Nikolaos P. Daskalakis[7], Jose Davila-Velderrain[17], Amy Deep[8], Chengyu Deng[3], Chris DiPietro[7], Stella Dracheva[1], Shiron Drusinsky[18], Ziheng Duan[19], Duc Duong[20], Cagatay Dursun[10], Nick Eagles[8], Jonathan Edelstien[1],Prashant S. Emani[10],John Fullard[1], Kiki Galani[21], Timur Galeev[10], Michael J. Gandal[11], Sophia Gaynor[12], Mark Gerstein[10], Daniel Geschwind[9], Kiran Girdhar[1], Fernando S. Goes[22], William Greenleaf[5], Jennifer Grundman[9], Qiuyu Guo[9], Chirag Gupta[23], Yoav Hadas[1], Joachim Hallmayer[5], Xikun Han[21], Vahram Haroutunian[1], Natalie Hawken[9], Chuan He[24], Ella Henry[10], Joo Heon Shin[8], Stephanie Hicks[8], Marcus Ho[5], Li-Lun Ho[21], Gabriel E. Hoffman[1], Yiling Huang[5], Louise Huuki[8], Ahyeon Hwang[19], Thomas Hyde[8], Artemis Iatrou[7], Fumitaka Inoue[3], Aarti Jajoo[7], Matthew Jensen[10], Lihua Jiang[5], Peng Jin[20], Ting Jin[23], Connor Jops[11], Alexandre Jourdon[10], Riki Kawaguchi[9], Manolis Kellis[21], Joel Kleinman[8], Steven P. Kleopoulos[1], Alex Kozlenkov[1], Arnold Kriegstein[3], Anshul Kundaje[5], Soumya Kundu[5], Cheyu Lee ,University California Irvine[19], Donghoon Lee[1], Junhao Li[13], Mingfeng Li[10], Xiao Lin[1], Shuang Liu[10], Jason Liu[10], Jianyin Liu[9], Chunyu Liu[16], Shuang Liu[23], Shaoke Lou[10], Jacob Loupe[14], Dan Lu[25], Shaojie Ma[10], Liang Ma[26], Michael Margolis[9], Jessica Mariani[10], Keri Martinowich[8], Kristen R. Maynard[8], Samantha Mazariegos[9], Ran Meng[10], Richard Meyers[14], Courtney Micallef[1], Tatiana Mikhailova[16], Guo-li Ming[11], Shahin Mohammadi[27], Emma Monte[5], Kelsey S. Montgomery[25], Jill E. Moore[28], Jennifer Moran[12], Eran Mukamel[13], Angus Nairn[10], Charles Nemeroff[29], Pengyu Ni[10], Scott Norton[10], Tomasz Nowakowski[3], Larsson Omberg[25], Stephanie C. Page[8], Saejeong Park[10], Ashok Patowary[9], Reenal Pattni[5], Geo Pertea[8], Mette A. Peters[25], Nishigandha Phalke[28], Dalila Pinto[1], Milos Pjanic[1], Sirisha Pochareddy[10], Katherine Pollard[18], Alex Pollen[3], Henry Pratt[28], Pawel F. Przytycki[18], Carolin Purmann[5], Zhaohui S. Qin[20], Ping-Ping Qu[5], Diana Quintero[9], Towfique Raj[1], Ananya S. Rajagopalan[10], Sarah Reach[1] , Thomas Reimonn[28], Kerry J. Ressler[7], Deanna Ross[4], Panagiotis Roussos[1], Joel Rozowsky[10], Misir Ruth[1], W. Brad Ruzicka[7], Stephan J. Sanders[30], Juliane M. Schneider[25], Soraya Scuderi[10], Robert Sebra[1], Nenad Sestan[10], Nicholas Seyfried[20], Zhiping Shao[1], Nicole Shedd[28], Annie W. Shieh[31], Mario Skarica[10], Clara Snijders[7], Hongjun Song[11], Matthew State[3], Jason Stein[32], Marilyn Steyert[3], Sivan Subburaju[7], Thomas Sudhof[5], Michael Synder[5], Ran Tao[8], Karen Therrien[1], Li-Huei Tsai[21], Alexander Urban[5], Flora M. Vaccarino[10], Harm van Bakel[1], Daniel Vo[11], Georgios Voloudakis[1], Brie Wamsley[9], Tao Wang[5], Sidney H. Wang[31], Daifeng Wang[23], Yifan Wang[2], Jonathan Warrell[10], Yu Wei[16], Annika Weimer[5], Daniel R. Weinberger[8], Cindy Wen[9], Zhiping Weng[28], Sean Whalen[18], Kevin White[33], A Jeremy. Willsey[3], Hyejung Won[32], Wing Wong[5], Hao Wu[20], Feinan Wu[10], Stefan Wuchty[34], Dennis Wylie[4], Siwei Xu[19], Chloe X. Yap[34], Biao Zeng[1], Pan Zhang[9], Chunling Zhang[16], Bin Zhang[1], Jing Zhang[19], Yanqiong Zhang[32], Xiao Zhou[10], Ryan Ziffra[3], Trisha M. Zintel[25]

[1]Icahn School of Medicine at Mount Sinai, New York, NY, USA. [2]Mayo Clinic Rochester, Rochester, MN, USA. [3]University of California, San Francisco, San Francisco, CA, USA. [4]The University of Texas at Austin, Austin, TX, USA. [5]Stanford University, Stanford, CA, USA. [6]University of Washington, Seattle, WA, USA. [7]McLean Hospital, Belmont, MA, USA. [8]Lieber Institute for Brain Development, Baltimore, MD, USA. [9]University of California, Los Angeles, Los Angeles, CA, USA. [10]Yale University, New Haven, CT, USA. [11]University of Pennsylvania, Philadelphia, PA, USA. [12]Tempus Labs, Inc., Chicago, IL, USA. [13]University of California, San

Diego, San Diego, CA, USA. [14]HudsonAlpha Institute for Biotechnology, Huntsville, AL, USA. [15]Duke University, Durham, NC, USA. [16]SUNY Upstate Medical University, Syracuse, NY, USA. [17]Human Technopole, Milan, Italy. [18]Gladstone Institutes, University of California, San Francisco, San Francisco, CA, USA. [19]University of California, Irvine, Irvine, CA, USA. [20]Emory University, Atlanta, GA, USA. [21]Massachusetts Institute of Technology, Cambridge, MA, USA. [22]Johns Hopkins University, Baltimore, MD, USA. [23]University of Wisconsin-Madison, Madison, WI, USA. [24]The University of Chicago, Chicago, IL, USA. [25]Sage Bionetworks, Seattle, WA, USA. [26]The University of Texas Health Science Center at San Antonio, San Antonio, TX, USA. [27]Broad Institute of MIT and Harvard, Cambridge, MA, USA. [28]University of Massachusetts Chan Medical School, Worcester, MA, USA. [29]The University of Texas at Austin Dell Medical School, Austin, MA, USA. [30]University of Oxford, Oxford, England, UK. [31]The University of Texas Health Science Center at Houston, Houston, TX, USA. [32]University of North Carolina at Chapel Hill, Chapel Hill, USA. [33]National University of Singapore, Singapore, Singapore. [34]University of Miami, Miami, FL, USA. University of Queensland, Queensland, NZ.
